# Functional classification of *GNAI1* disorder variants in *C. elegans* uncovers conserved and cell-specific mechanisms of dysfunction

**DOI:** 10.1101/2025.08.18.670960

**Authors:** Rehab Salama, Eric Peet, Logan Morrione, Sarah Durant, Maxwell Seager, Madison Rennie, Suzanne Scarlata, Inna Nechipurenko

## Abstract

Heterotrimeric G proteins transduce signals from G protein coupled receptors, which mediate key aspects of neuronal development and function. Mutations in the *GNAI1* gene, which encodes Gαi1, cause a disorder characterized by developmental delay, intellectual disability, hypotonia, and epilepsy. However, the mechanistic basis for this disorder remains unknown. Here, we show that *GNAI1* is required for ciliogenesis in human cells and use *C. elegans* as a whole-organism model to determine the functional impact of seven *GNAI1*-disorder patient variants. Using CRISPR-Cas9 editing in combination with robust cellular (cilia morphology) and behavioral (chemotaxis) assays, we find that *T48I*, *K272R*, *A328P*, and *V334E* orthologous variants impact both cilia assembly and function in AWC neurons, *M88V* and *I321T* have no impact on either phenotype, and *D175V* exerts neuron-specific effects on cilia-dependent sensory behaviors. Finally, we validate in human ciliated cell lines that *D173V*, *K270R*, and *A326P GNAI1* variants disrupt ciliary localization of the encoded human Gαi1 proteins similarly to their corresponding orthologous substitutions in the *C. elegans* ODR-3 *(D175V*, *K272R*, and *A328P*). Overall, our findings determine the *in vivo* effects of orthologous *GNAI1* variants and contribute to mechanistic understanding of *GNAI1* disorder pathogenesis as well as neuron-specific roles of ODR-3 in sensory biology.

**ARTICLE SUMMARY:** Gα subunits of heterotrimeric G proteins transduce signaling from G protein coupled receptors and play important roles in cell communication and complex behaviors. Mutations in the *GNAI1* gene, which encodes Gαi1 protein, have been recently linked to a neurodevelopmental disorder; however, it remains unknown how *GNAI1* patient mutations disrupt neuronal development or function to manifest in disease. We demonstrate that *GNAI1* is required for ciliogenesis and use *C. elegans* as a whole-animal model in combination with human cells to identify cell-specific and conserved mechanisms of Gα dysfunction.

## INTRODUCTION

Heterotrimeric G proteins are key transducers of G protein coupled receptor (GPCR) signaling, which plays key roles in neuronal development, communication, and behavior (Betke et al., 2012; Kurabayashi et al., 2013; Wettschureck & Offermanns, 2005). In the canonical GPCR cascade, the activated receptor acts as a guanine-nucleotide exchange factor (GEF) to exchange guanosine diphosphate (GDP) on the Gα subunit of the Gαβγ trimer for guanosine triphosphate (GTP), thus stimulating dissociation of Gα-GTP from Gβγ and allowing activation of their respective downstream effectors (Dror et al., 2015; Gilman, 1987; Pierce et al., 2002).

Many GPCRs, G proteins, and their downstream effectors localize to primary cilia – specialized cellular compartments that house molecular machinery of all major signaling pathways (Anvarian et al., 2019; Hilgendorf et al., 2016). Primary cilia mediate many aspects of neuronal biology in the developing and mature brain including cell fate specification, proliferation, migration, axon guidance, dendrite morphogenesis, and neuronal excitability (Hasenpusch-Theil & Theil, 2021; Jurisch-Yaksi et al., 2024; Stoufflet & Caille, 2022; Suciu & Caspary, 2021). In humans, cilia dysfunction is associated with a spectrum of developmental disorders called ciliopathies (Hildebrandt et al., 2011; Reiter & Leroux, 2017). Ciliopathy patients commonly exhibit neurological symptoms that range in severity and include structural brain abnormalities, intellectual disability, motor deficits, and epilepsy (Guemez-Gamboa et al., 2014; Lee & Gleeson, 2011; Valente et al., 2014), further highlighting the critical importance of primary cilia in the nervous system. In addition to classic ciliopathies, cilia defects are being increasingly reported in neurodevelopmental, neurodegenerative, and psychiatric disorders suggesting that cilia dysfunction may be a shared feature of many neurological conditions (Jurisch-Yaksi et al., 2024; Karalis et al., 2022). However, the mechanisms by which cilia dysfunction contributes to neurological phenotypes observed in these disorders or those by which risk genes for these disorders impact cilia biology remain largely unknown.

Recent studies identified >15 *de novo* mutations in the *GNAI1* gene, which encodes the Gαi1 subunit of heterotrimeric G (αβγ) proteins, in individuals with a novel neurodevelopmental disorder (NDD) henceforth referred to as ‘*GNAI1* disorder’ (Muir et al., 2021). Most affected individuals exhibited developmental delay, intellectual disability, hypotonia, and epilepsy – neurological symptoms that are also commonly associated with classic ciliopathies. Although the mechanistic basis for *GNAI1* disorder remains unknown, prior studies have described several neuronal functions for Gαi1. In the canonical GPCR cascade, Gαi1 is responsible for decreasing cAMP levels upon receptor activation by inhibiting adenylyl cyclases (Wettschureck & Offermanns, 2005). Acute *GNAI1* knockdown in the mouse embryonic cortex disrupted multiple aspects of cortical development including progenitor proliferation, neuronal migration, and dendritogenesis (Hamada et al., 2021). Additionally, genetic ablation of *GNAI1* has been reported to increase adenylyl cyclase activity and impair hippocampus-dependent long-term memory in mice (Pineda et al., 2004). While the cellular and molecular mechanisms that underlie neuronal functions of Gαi1 remain to be fully elucidated, these studies demonstrate that Gαi1 plays critical roles both in the developing and mature brain and underscore the importance of understanding the functional effects of *GNAI1* patient variants in relevant cellular and developmental contexts *in vivo*.

*C. elegans* constitutes a powerful whole-organism platform for rapidly evaluating phenotypic consequences of NDD-associated missense variants. Several recent studies successfully deployed *C. elegans* to functionally classify *de novo* variants in risk genes for developmental disorders that include *GNAO1* encephalopathy (Wang et al., 2022), autism spectrum disorder (Wong et al., 2019), and ciliopathies (Lange et al., 2022; Lange et al., 2021). Here, we use *C. elegans* ODR-3, which belongs to the Gαi/o class of G proteins, as a model to evaluate the functional impacts of seven NDD-associated *GNAI1* variants. Using *C. elegans* ODR-3 for this purpose offers several key advantages. First, all missense variants selected for analysis alter amino acids that are identical between Gαi1 and ODR-3 and map to the functional motifs conserved across Gα proteins. Second, *odr-3* loss-of-function (lf) and gain-of-function (gf) mutations result in robust and easily quantifiable cellular phenotypes – defects in cilia morphology in AWA and AWC olfactory neurons (Campagna et al., 2023; Lans et al., 2004; Roayaie et al., 1998). Finally, *odr-3* function is required for a panel of chemosensory behaviors including avoidance of high-osmolarity solutions detected by ASH and chemotaxis toward attractive odorants detected by AWA and AWC neurons (Bargmann et al., 1993; Roayaie et al., 1998). As a result, a complementary approach combining robust cellular and behavioral assays can be used to rapidly identify gene variants that are functionally consequential *in vivo*.

We find that five of the examined orthologous variants exhibited a range of defects in AWC cilia morphology and/or AWC-mediated attraction to benzaldehyde, while two variants appeared wild-type in both assays. Interestingly, one examined variant had cell-specific effects on neuronal function - a marked deficit in ASH-mediated glycerol avoidance and unaltered AWC-mediated chemotaxis toward benzaldehyde. Finally, we confirm that the functional impact of three orthologous variants that severely compromised localization of the encoded ODR-3 proteins to AWC cilia is conserved in human cells.

## MATERIALS AND METHODS

### *C. elegans* strains and maintenance

All *C. elegans* strains were maintained at 15°C or 20°C on nematode growth medium (NGM) agar plates seeded with *Escherichia coli* OP50 (Brenner, 1974). Standard genetic techniques were used to generate all strains. PCR and/or Sanger sequencing was used to confirm genotypes of all mutant strains. Transgenic *C. elegans* carrying multicopy extrachromosomal arrays were generated by germline transformation. To generate transgenic strains, experimental plasmids were microinjected at 5 – 10 ng/μl together with either *unc-122Δ*p*::gfp* or *unc-122Δ*p*::dsred* co-injection markers injected at 30 ng/μL and 40 ng/μL, respectively. Wild-type and all mutant TagRFP-tagged *odr-3* constructs were injected at identical concentrations. Two or more independently generated lines were examined for all extrachromosomal transgenes, and the same extrachromosomal array was examined in wild-type and mutant backgrounds that were being directly compared. A list of *C. elegans* strains used in this study is provided in Supplementary Table 1.

### Chemotaxis behavioral assays

Population chemotaxis assays were performed on 10-cm round petri plates as previously described (Bargmann et al., 1993; Campagna et al., 2023). Briefly, 1 mL of benzaldehyde (Sigma) diluted in ethanol at 1:200 and 1 mL of ethanol (diluent control) were placed along with 1 mL of 1M sodium azide anesthetic at opposite ends of the assay plate. Young adult hermaphrodites were washed off their growth plates with cholesterol-free S-basal, washed twice with S-basal, once with Milli-Q water, and transferred to assay plates. After one hour, chemotaxis index was calculated as follows:

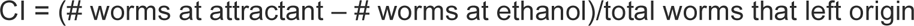

All assays were performed on three or more days with at least two technical replicates per genotype per day. Wild-type and *odr-3(lf)* controls were included with experimental genotypes on all days of the assay.

### Osmotic avoidance assays

Osmotic avoidance assays were performed as previously described (Cornils et al., 2016; Culotti & Russell, 1978). Briefly, 4-6 one-day old hermaphrodites were transferred from a growth plate onto an NGM plate without food for 2 minutes to remove residual *E. coli* OP50. The same animals were subsequently placed inside a ring (∼3/8^th^ inch in diameter) of 8M glycerol colored with bromophenol blue (USB). After 2 minutes, the number of animals that remained inside the glycerol ring was counted. All assays were repeated on five or more separate days with at least three technical replicates per genotype per day. Wild-type and *odr-3(lf)* controls were included with experimental genotypes on all days of the assay.

### CRISPR-Cas9 gene editing

Cas9 protein, crRNAs, and tracrRNA were purchased from Integrated DNA Technologies (IDT). All missense mutations and insertion of the split-wrmScarlet (wrmScarlet_11_) tag were confirmed by Sanger sequencing.

#### Generation of missense odr-3 variants

gene editing was carried out as described in (Dokshin et al., 2018). Briefly, donor oligonucleotides carrying the desired missense mutations, silent mutations within the crRNA target sequence and/or PAM site to prevent repetitive cutting, and 36-40 bp homology arms were ordered from IDT (see Supplementary Fig. 4 for detailed sequence information). The donor template (25 ng/mL), crRNA (56 ng/mL), tracrRNA (100 ng/mL), and Cas9 protein (250 ng/mL) were co-injected with *unc-122D*p*::dsred* or *unc-122D*p*::gfp* co-injection markers, and F1 progeny expressing the co-injection marker were isolated and genotyped for the presence of the edit. F2 individuals homozygous for the desired mutation were subsequently isolated from heterozygous F1 parents. All mutants generated via CRISRP-Cas9 editing were sequenced by Sanger sequencing and outcrossed at least two times to mitigate potential off-target effects.

#### Generation of wrmScarlet_11_ knock-in strain

To generate the split-wrmScarlet (*wrmScarlet_11_*) allele *odr-3(nch013),* a single-stranded donor oligonucleotide containing the *wrmScarlet_11_* fragment (Goudeau et al., 2021), linkers, and homology arms was synthesized by IDT. *wrmScarlet_11_* sequence was inserted between codons encoding Gly118 and Glu119 residues of ODR-3 (Campagna et al., 2023).

crRNA: 5’ GTACAGGAAAATGGAGAAGA 3’

donor oligonucleotide: 5’ AAGAAGCAGAAAAGGCAATAGTTATGAAAGTACAGGAAAAcGGcGAgGAAGGAAGTGGAGGAGGAGGAAGTTACACCGTCGTCGAGCAATACGAGAAGTCCGTCGCCCGTCACTGCACCGGAGGAATGGATGAGTTATACAAGAGTGGAGGAGGAGGAAGTGAAGCACTGACAGAAGAAGTTTCGAAAGCAATTCAATCG 3’

### Molecular biology

#### odr-3 mutant constructs

expression of *odr-3* transgenes in AWC and ASH was achieved by sub-cloning *odr-3* cDNA downstream from 0.7 kb of *ceh-36* (*ceh-36*Δp) (Kim et al., 2010) and ∼3 kb of *sra-6* (*sra-6*p) regulatory sequences, respectively. TagRFP tag was inserted between Gly118 and Glu119 residues of ODR-3 as previously described (Campagna et al., 2023). Patient mutations were introduced into plasmids carrying wild-type *tagrfp*-tagged *odr-3* cDNA downstream from cell-specific promoters by site-directed mutagenesis using QuikChange Lightning kit (Agilent Technologies). All constructs were verified by whole-plasmid sequencing (Plasmidsaurus).

#### GNAI1 constructs

wild-type coding sequence of human *GNAI1* (NM 002069) cloned into *pcDNA3.1+* (Invitrogen) mammalian expression vector was purchased from Bloomsburg University Foundation cDNA Resource Center (Catalog # GNAI100000). *eGFP* coding sequence with flanking DNA segments encoding GlyThr (5’ linker) and GlySer (3’ linker) was inserted between Leu91 and Lys92 of *GNAI1* coding sequence in *pcDNA3.1+* vector using NEBuilder HiFi DNA assembly (NEB) to make *eGFP*-tagged *GNAI1* construct. Missense mutations corresponding to *D173V*, *K270R*, and *A326P* substitutions were introduced into the *pcDNA3.1+* plasmid containing *eGFP*-tagged *GNAI1* via site-directed mutagenesis using QuikChange Lightning kit (Agilent Technologies). All constructs were verified by full-plasmid sequencing (Plasmidsaurus). A list of plasmids used in this work is provided in Supplementary Table 2.

#### qPCR

total RNA was extracted from siControl and si*GNAI1*-treated RPE-1 and HEK293T cells using RNeasy kit (Qiagen) per manufacturer’s instructions. RNA samples were reverse-transcribed (ZymoScript One-Step RT-qPCR Kit), and *GNAI1* expression (relative to *RPL11* control) was quantified by real-time PCR carried out on Applied Biosystems QuantStudio 6 Pro system using the 2^-ΔΔCt^ method. Primer pairs used in this study are below:

*RPL11*: 5’ GTTGGGGAGAGTGGAGACAG 3’ / 5’ TGCCAAAGGATCTGACAGTG 3’

*GNAI1*: 5’ CCCGAGAGTACCAGCTTAATG 3’ / 5’ CATCTTGTTGAGTCGGGATGTA 3’

### Cell culture and transfections

All cell lines were maintained at 37°C with 5% CO_2_ and tested monthly for mycoplasma using mycoplasma PCR detection kit (ABM). Human telomerase-immortalized retinal pigment epithelial cells (hTERT RPE-1 (Nechipurenko et al., 2016)) were cultured in DMEM/F-12 (1:1) supplemented with 10% fetal bovine serum (FBS) and 1X antibiotic-antimycotic (Gibco). HEK293T cells (Abcam) were grown in DMEM high-glucose supplemented with Glutamax (Gibco), 10% FBS, and 1X antibiotic-antimycotic (Gibco).

#### RNAi

For immunofluorescence analysis, RPE-1 and HEK293T cells were plated on 12-mm glass pre-treated or Poly-D-Lysine-coated coverslips (Neuvitro), respectively, in 24-well plates at 30,000 cells per well in antibiotic-free complete-growth medium. For qPCR analysis, cells were plated without coverslips in 12-well plates at 60,000 cells per well. Synthetic small interfering RNA oligonucleotides (siRNAs) targeting non-overlapping *GNAI1* sequences or non-targeting control siRNA were transfected as previously described (Nechipurenko et al., 2016) using Lipofectamine RNAiMax (Invitrogen). The target sequences and sources of siRNAs used in this study are provided below:

si*GNAI1* #1 (s5872, Ambion): GAAUUGUUGAAACCCAUUU

si*GNAI1* #2 (J-010404-07-0002, Dharmacon): CAAAUUACAUCCCGACUCA

siControl (D-001810-01-05, Dharmacon): UGGUUUACAUGUCGACUAA

#### Transfection of GNAI1 plasmids

HEK293T cells were plated on 12-mm glass Poly-D-Lysine-coated coverslips (Neuvitro) in 24-well plates at 30,000 cells per well in antibiotic-free complete-growth medium. Upon reaching 50-60% confluence, the plated cells were transfected with 500 ng of pcDNA3.1+ vector containing eGFP-tagged *GNAI1^WT^*, *GNAI1^D173V^*, *GNAI1^K270R^*, or *GNAI1^A326P^* using Lipofectamine 3000 (Invitrogen) per manufacturer’s instructions. After a three-day incubation period, the transfection medium was replaced with serum-free medium to induce ciliation. Cells were serum-starved for 48 hrs prior to fixation and analysis.

### Immunostaining

RPE-1 and HEK293T cells were fixed in 4% paraformaldehyde for 12 minutes followed by permeabilization in 0.2% Triton X-100 (MP Biomedicals) for 10 minutes at room temperature (RT). Fixed cells were blocked with 5% bovine serum albumin in phosphate buffered saline (PBS) with 0.2% Triton X-100 (PBS-T) for 1 hour at RT or at 4°C overnight, followed by incubation in primary antibodies for 1.5 hours at RT or at 4°C overnight. The following antibodies were used in immunofluorescence experiments: anti-γ-tubulin (clone 8D11, 1:500, Biorbyt, batch # T1578), anti-ARL13B (clone N295B/66, 1:10, Developmental Studies Hybridoma Bank), anti-acetylated α-tubulin (clone 6-11B-1, 1:500, Sigma, batch # T7451), anti-GFP (catalog # GFP-1010, 1:500, Aves Laboratories, batch # GFP917979), and anti-GM130/GOLGA2 (catalog # HPA021178, 1:500, Sigma, batch # A105115). Species-specific fluorescent secondary antibodies were obtained from Jackson ImmunoResearch laboratories and used at 1:500 dilution in PBS-T. DAPI (1:1000, ThermoFisher) was used to stain DNA and applied together with secondary antibodies for 1.5 hours at RT or at 4°C overnight.

### Microscopy

#### C. elegans

Synchronized one-day old adult hermaphrodites were anesthetized in 10mM tetramisole hydrochloride (MP Biomedicals) and placed on a 10% agarose pad mounted on a microscope slide. Animals were imaged on an upright THUNDER Imager 3D Tissue (Leica). Complete z-stacks of AWC and ASH sensory endings that included distal dendrite and cilia were acquired at 0.22-μm intervals with a K5 sCMOS camera (Leica) in Leica Application Suite X software using an HC Plan Apochromat 63X NA 1.4–0.60 oil immersion objective. To image one-day-old split-wrmScarlet transgenic animals, single-plane snapshots were acquired on an inverted Nikon Ti-E microscope with Yokogawa CSU-X1 spinning disk confocal head using 60X NA 1.40 oil immersion objective.

#### RPE-1 and HEK293T cells

Fixed and stained RPE-1 and HEK293T cells were mounted in ProLong Diamond antifade mountant (Invitrogen) and imaged on an inverted Nikon Ti-E microscope with Yokogawa CSU-X1 spinning disk confocal head using a 60X NA 1.40 oil immersion objective. Complete z-stacks were acquired at 0.25-μm intervals with an ORCA-Fusion BT Digital CMOS camera (Hamamatsu) in MetaMorph 7 software (Molecular devices).

### Image analysis

Analysis of all fluorescence microscopy images was performed in Fiji/Image J (National Institute of Health, Bethesda, MD). All data were quantified from images collected on at least two independent days. For experiments examining localization of TagRFP-tagged ODR-3 variants and eGFP-tagged Gαi1 variants, all genotypes/conditions being compared directly were imaged at identical settings.

#### ODR-3::TagRFP fluorescence intensity (AWC and ASH)

TagRFP fluorescence intensity inside AWC cilia or PCMC in the distal dendrite was measured by outlining AWC cilia or PCMC using the freehand selection tool and measuring mean intensity inside the resultant ROIs. The relative fluorescence intensity was plotted as a ratio of mean TagRFP intensity inside the AWC cilium over mean TagRFP intensity inside the PCMC of the same neuron. TagRFP fluorescence intensity inside ASH cilium was measured by drawing a line segment along the center of the ASH cilium (from base to tip) and measuring mean intensity along the line. TagRFP intensity inside the PCMC of ASH was measured as described above for AWC, and the relative intensity was calculated and plotted as a ratio of intensity inside the ASH cilium over intensity in the PCMC of the same neuron.

#### Cilia area (AWC)

The area of AWC cilia was measured as previously described (Campagna et al., 2023). Briefly, the cilia were outlined using the freehand selection tool, and the area enclosed by the ROI was measured from maximum-intensity projections that contained AWC cilia in their entirety.

#### Gαi1::eGFP relative fluorescence intensity

Line segments were drawn from base to tip, along the center of the cilia, visualized using anti-ARL13B antibody, and mean intensity along the line was measured in the eGFP channel in transfected HEK293T cells. Another straight line was drawn across the Golgi visualized with anti-GOLGA2 antibody, and mean intensity along the line was measured in the eGFP channel. The relative Gαi1::eGFP intensity was calculated and plotted as a ratio of eGFP intensity inside the cilium over eGFP intensity in the Golgi of the same transfected cell.

### Statistical analysis

Prism 10 software (GraphPad, San Diego, CA) was used to perform all statistical analyses and generate bar graphs and scatter plots. The D’Agostino-Pearson test was used to determine whether the data were normally distributed. Details on the numbers of analyzed animals and cells, statistical tests, and multiple comparison correction (when applicable) are presented in figure legends.

## RESULTS

### *GNAI1* is required for ciliogenesis in mammalian cells

Given the overlap in clinical features observed in classic ciliopathies and *GNAI1* disorder (Neurodevelopmental disorder with hypotonia, impaired speech, and behavioral abnormalities; OMIM # 619854), we first wondered whether *GNAI1* may play a role in cilia assembly. To address this question, we knocked down *GNAI1* in human embryonic kidney (HEK293T) and immortalized retinal pigment epithelial (hTERT RPE-1) cells – two commonly used cell culture models in cilia research. Roughly 80% of RPE-1 cells transfected with non-targeting control siRNA (*si*Control) possessed a primary cilium labeled with anti-ARL13B antibody; however, only 34% of cells transfected with si*GNAI1* (si*GNAI1 #1*) that targets both *GNAI1* reference transcripts (NM 001256414.1 and NM 002069.5) were ciliated (Fig. 1a and c), suggesting that *GNAI1* is required for ciliogenesis in RPE-1 cells. Importantly, RPE-1 cells transfected with the second siRNA that targets a distinct non-overlapping region of *GNAI1* (si*GNAI1* #2) resulted in a similar knockdown efficiency and ciliation defect (Supplementary Fig. 1 a and b), suggesting that the observed cilia phenotype is likely *GNAI1*-dependent. To confirm that reduction in cilia number observed upon *GNAI1* KD is not simply due to defective ciliary trafficking of ARL13B, we stained *GNAI1* KD RPE-1 cells with an antibody against acetylated tubulin, which labels the axoneme, and noted a comparable reduction in number of ciliated cells (Supplementary Fig. 1 c and d).

**Fig. 1.**
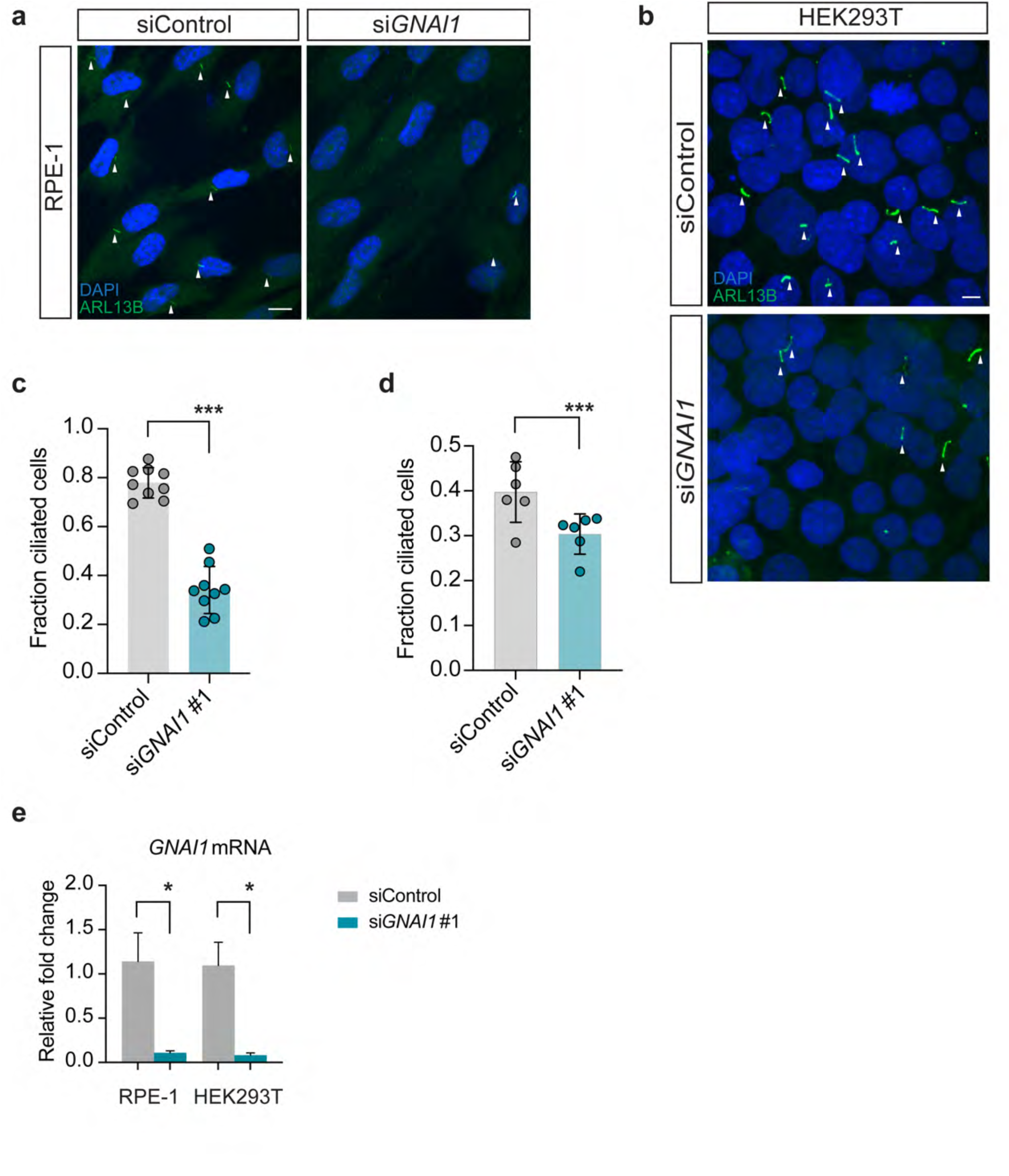
*GNAI1* knockdown impairs ciliogenesis in human cells. a – b) Immunofluorescence images of fixed RPE-1 (a) and HEK293T (b) cells transfected with the indicated siRNAs. White arrowheads mark cilia labeled with anti-ARL13B antibody. Scale: 10 μm. c – d) Quantification of ciliation in RPE-1 (c) and HEK293T (d) cells transfected with the indicated siRNAs. Each data point represents one replicate. Total number of cells: siControl (n=858 RPE-1, n=4460 HEK293T), si*GNAI1* #1 (n=518 RPE-1, n=4600 HEK293T). Means ± SD are indicated by shaded and vertical bars, respectively. *** indicates different from control at p<0.001 (Fisher’s exact test). e) Relative levels of *GNAI1* mRNA in RPE-1 and HEK293T cells treated with the indicated siRNAs. Summary data represent > 3 replicates. Error bars are SEM. * indicates different from control at p<0.05 (Mann-Whitney test).

To test if *GNAI1* function is broadly required for ciliogensis in human cells, we next turned to HEK293T cells. Similarly to RPE-1 cells, *GNAI1* KD in HEK293T cells caused a significant reduction in ciliogenesis (Fig. 1b and d). However, the overall impact of *GNAI1* KD on ciliation of HEK293T cells appeared to be qualitatively milder relative to RPE-1 cells (compare Fig. 1c and Fig. 1d), despite similar *GNAI1* KD efficiency in both cell lines (Fig. 1e), indicating that *GNAI1* may differentially contribute to cilia assembly in different cell types.

### Selection of *GNAI1* disorder variants for *in vivo* functional analysis

Sequence analysis determined that *C. elegans* ODR-3 exhibits 49% amino acid sequence identity and 66% similarity to human Gαi1 (100% length, EMBOSS Needle; BLAST e-value 6×10^-127^; Supplementary Fig. 2). The majority of *GNAI1* disorder missense variants impact residues in the protein domains conserved across all Gα proteins (Muir et al., 2021). Therefore, we reasoned that we could leverage the well defined genetics of *odr-3* together with robust cellular and behavioral phenotypes displayed by *odr-3(lf)* and *odr-3(gf)* mutants to determine functional impacts of select *GNAI1* patient variants *in vivo*. To this end, we chose to focus on seven missense *GNAI1* alleles that affect conserved, identical amino acids in ODR-3 (Fig. 2a and b; Supplementary Fig. 2). Four mutations (*GNAI1*: *T48I*, *D173V*, *K270R*, and *A326P*) map to structurally conserved motifs called G boxes that mediate binding and hydrolysis of guanine nucleotides (Luo et al., 2022; Noel et al., 1993; Sprang, 1997), one mutation (*GNAI1*: *V332E*) impacts a residue in the C-terminal alpha-helix, which participates in Gα interactions with GPCRs (Masuho et al., 2023; Oldham et al., 2006) and cytoplasmic GEF RIC-8 (Kant et al., 2016; Seven et al., 2020; Zeng et al., 2019), and two mutations (*GNAI1*: *M88V* and *I319T*) are in the conserved residues outside of known functional domains (Fig. 2a and b).

**Fig. 2.**
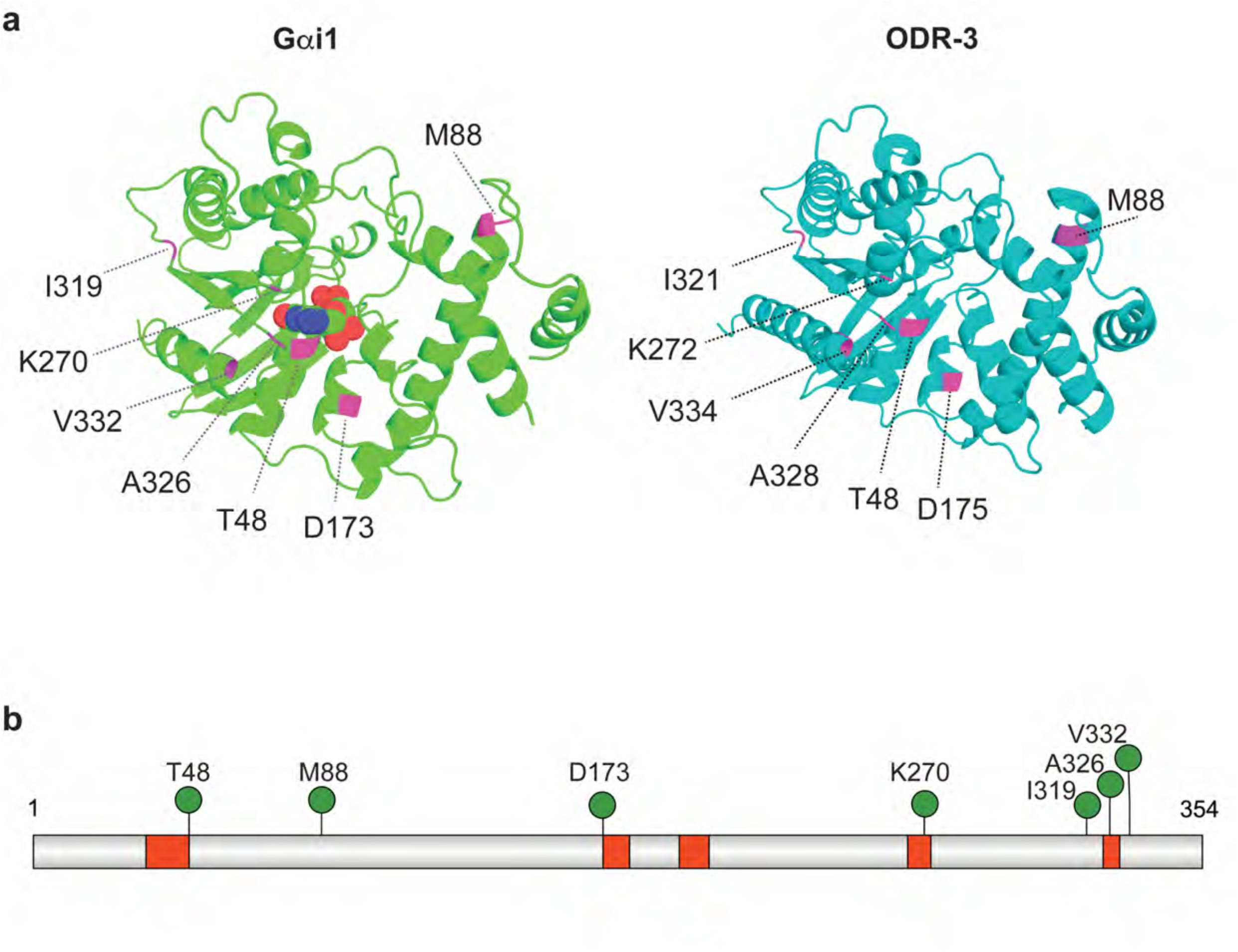
Distribution of patient variants selected for analysis. a) *Left*: 3D structure of Gαi1 bound to GDP (PDB accession: 2G83). Carbon, oxygen, nitrogen, and phosphorous atoms of GDP are shown in green, red, blue, and orange, respectively. *Right*: AlphaFold-predicted structure of *C. elegans* ODR-3 (Varadi et al., 2022; Varadi et al., 2024). Positions of missense patient variants that were selected for analysis are shown in magenta in both protein models. b) Schematic of human Gαi1 showing positions of *GNAI1* disorder-associated variants (green pins). Guanine nucleotide-binding motifs (G1 – G5 boxes) are shown in red.

### *GNAI1* disorder-associated orthologous variants differentially impact ODR-3 ciliary localization

Previous studies used overexpressed ODR-3 transgenes and immunofluorescence against endogenous ODR-3 to demonstrate that wild-type ODR-3 (ODR-3^WT^) is normally enriched in the cilia of AWC sensory neurons (Roayaie et al., 1998). First, we wanted to determine whether any of the selected variants affect sub-cellular localization of ODR-3. To address this question, we first tagged the *odr-3* locus with split-wrmScarlet reporter (*wrmScarlet_11_*) (Goudeau et al., 2021) using CRISPR-Cas9 genome editing. To visualize endogenous ODR-3 in AWC cilia, we reconstituted fluorescence via expression of the wrmScarlet_1-10_ fragment in AWC neurons under the *ceh-36Δ* promoter (Kim et al., 2010). Consistent with published studies, ODR-3^WT^::split-wrmScarlet was enriched inside AWC cilia (n=25 animals, Supplementary Fig. 3). However, the reconstituted wrmScarlet was photobleached nearly instantaneously upon sample illumination with the laser light making it impossible to acquire multiple *z*-slices necessary for in-depth quantitative analysis of ODR-3 subcellular localization.

To bypass the photobleaching issue, we next expressed TagRFP-tagged wild-type or mutant ODR-3 variants from transgenes in AWC neurons of wild-type animals and examined cilia localization of the encoded proteins. Similarly to the endogenous ODR-3, ODR-3^WT^::TagRFP expressed from a multicopy transgene was enriched in AWC cilia (Campagna et al., 2023) (Fig. 3a). We noted no significant changes in sub-cellular localization of TagRFP-tagged ODR-3^I321T^ or ODR-3^V334E^ variants (Fig. 3a and b). Interestingly, we observed two distinct patterns of mislocalization amongst the remaining ODR-3 protein variants. Like ODR-3^WT^, ODR-3^T48I^ and ODR-3^M88V^ were present inside the AWC cilium. However, unlike the WT protein, these variants were also detected in an ectopic pool in the periciliary membrane compartment (PCMC) of the distal dendrite (Fig. 3a and b). Finally, ODR-3^D175V^, ODR-3^K272R^, and ODR-3^A328P^ exhibited the most striking defects in sub-cellular localization – these mutant proteins were largely excluded from the AWC cilium and instead accumulated in the PCMC (Fig. 3a and b).

**Fig. 3.**
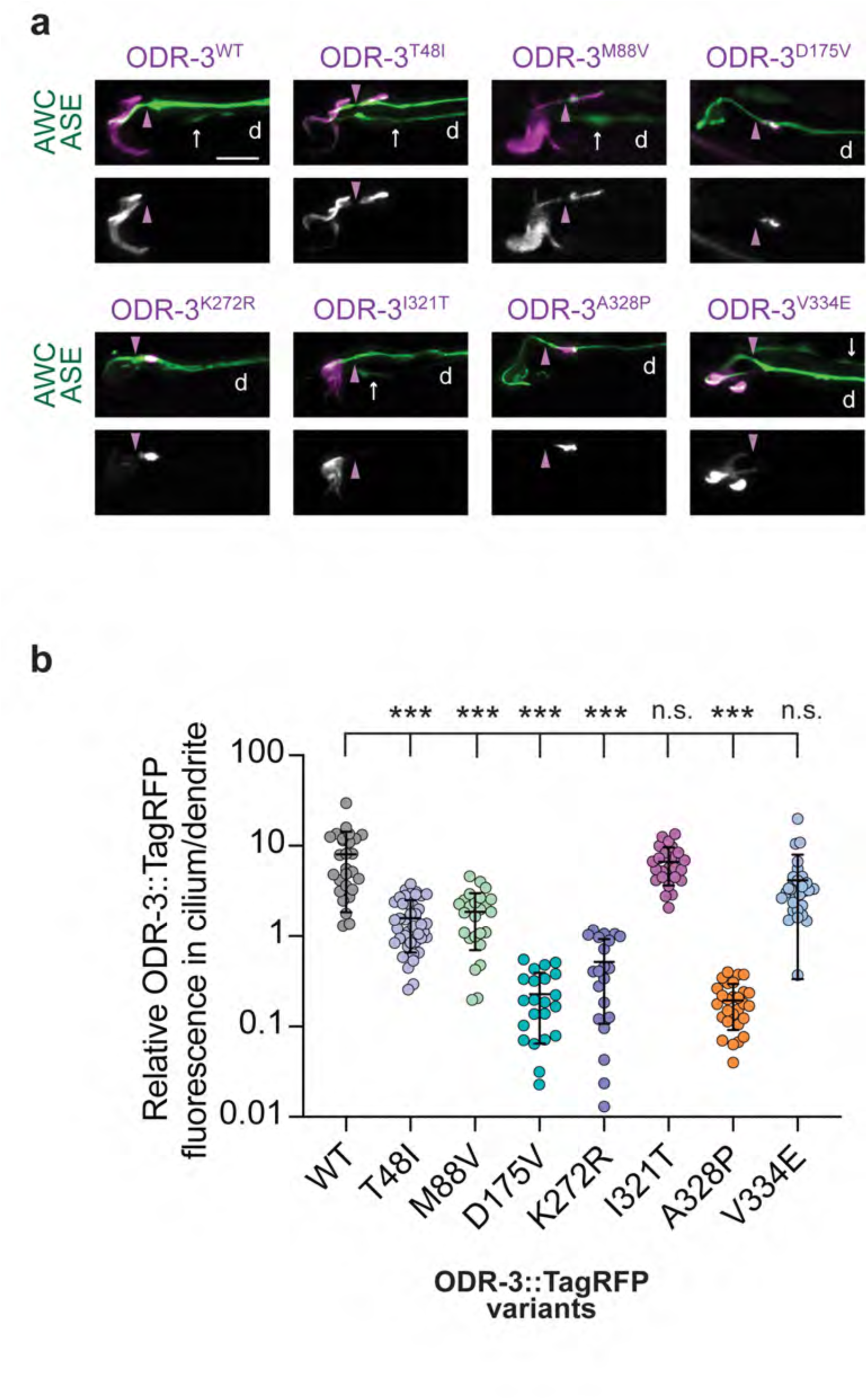
Orthologous patient mutations in the *odr-3* gene differentially affect subcellular localization of the encoded proteins. a) Representative images showing localization of the indicated TagRFP-tagged ODR-3 variants in AWC neurons. AWC neurons were visualized via expression of *ceh-36*p::GFP, which also labels ASE neurons (white arrows). Purple arrowheads mark cilia base. d: dendrite. Anterior is at left. Scale: 5 μm. b) Quantification of relative TagRFP fluorescence inside the AWC cilium vs distal dendrite for the indicated TagRFP-tagged ODR-3 variants. Total number of AWC neurons analyzed for each variant: WT (n=27), T48I (n=42), M88V (n=26), D175V (n=22), K272R (n=21), I321T (n=26), A328P (n=28), and V334E (n=29). Means ± SD are indicated by horizontal and vertical bars, respectively. *** indicates different from WT at p<0.001; n.s.: not significant (Kruskal-Wallis with Dunn’s multiple comparisons test).

To rule out the possibility that dendritic mislocalization observed with the mutant variants was an artifact of transgene overexpression, we introduced *A328P* mutation into the *odr-3::split-wrmScarlet* background. The endogenously tagged ODR-3^A328P^::split-wrmScarlet exhibited cilia localization defects that were qualitatively similar to those observed in transgenic animals overexpressing ODR-3^A328P^::TagRFP in AWC neurons (Supplementary Fig. 3). Specifically, ODR-3^A328P^::split-wrmScarlet was present throughout the AWC dendrite in all examined animals (n=30) and prominently accumulated at the cilia base in 28 out of 30 analyzed animals (Supplementary Fig. 3). Collectively, these data indicate that *T48I*, *M88V*, *D175V*, *K272R*, and *A328P* variants affect ciliary trafficking of the encoded ODR-3 proteins, with the latter three mutations having the most profound impact on ODR-3 ciliary localization.

### The transition zone contributes to ciliary exclusion of the ODR-3^A328P^ protein

Protein trafficking in and out of primary cilia is tightly controlled by the transition zone (TZ) – a selective diffusion barrier at the cilia base (Garcia-Gonzalo & Reiter, 2017). One possible explanation for the exclusion of ODR-3^A328P^ from the AWC cilium with a concomitant accumulation in the distal dendrite is that *A328P* mutation impedes ODR-3 transport into the cilium across the TZ. If this hypothesis were true, we would expect to see an increase in intra-ciliary levels of ODR-3^A328P^ protein in animals with disrupted TZ architecture compared to wild type. To test this hypothesis, we examined ODR-3^A328P^ localization in *mks-5(tm3100)* mutants that have compromised TZ integrity (Jensen et al., 2015). In line with our hypothesis, we noted an increase in ODR-3^A328P^::TagRFP levels inside the cilium relative to distal dendrite in *mks-5(tm3100)* mutants compared to ODR-3^A328P^::TagRFP levels in the wild-type background (Fig. 4a and b). Notably, when we examined localization of ODR-3^WT^::TagRFP in *mks-5(tm3100)* mutants, we noticed that in addition to cilia, ODR-3^WT^::TagRFP was now present in the distal dendrite, consistent with the previously published role for the TZ in preventing leakage of membrane-associated ciliary proteins into the dendrite (Cevik et al., 2013). Collectively, our results suggest that the TZ plays an important part in regulating ODR-3 ciliary entry and retention, and that the *A328P* variant likely impacts ODR-3 trafficking across the TZ.

**Fig. 4.**
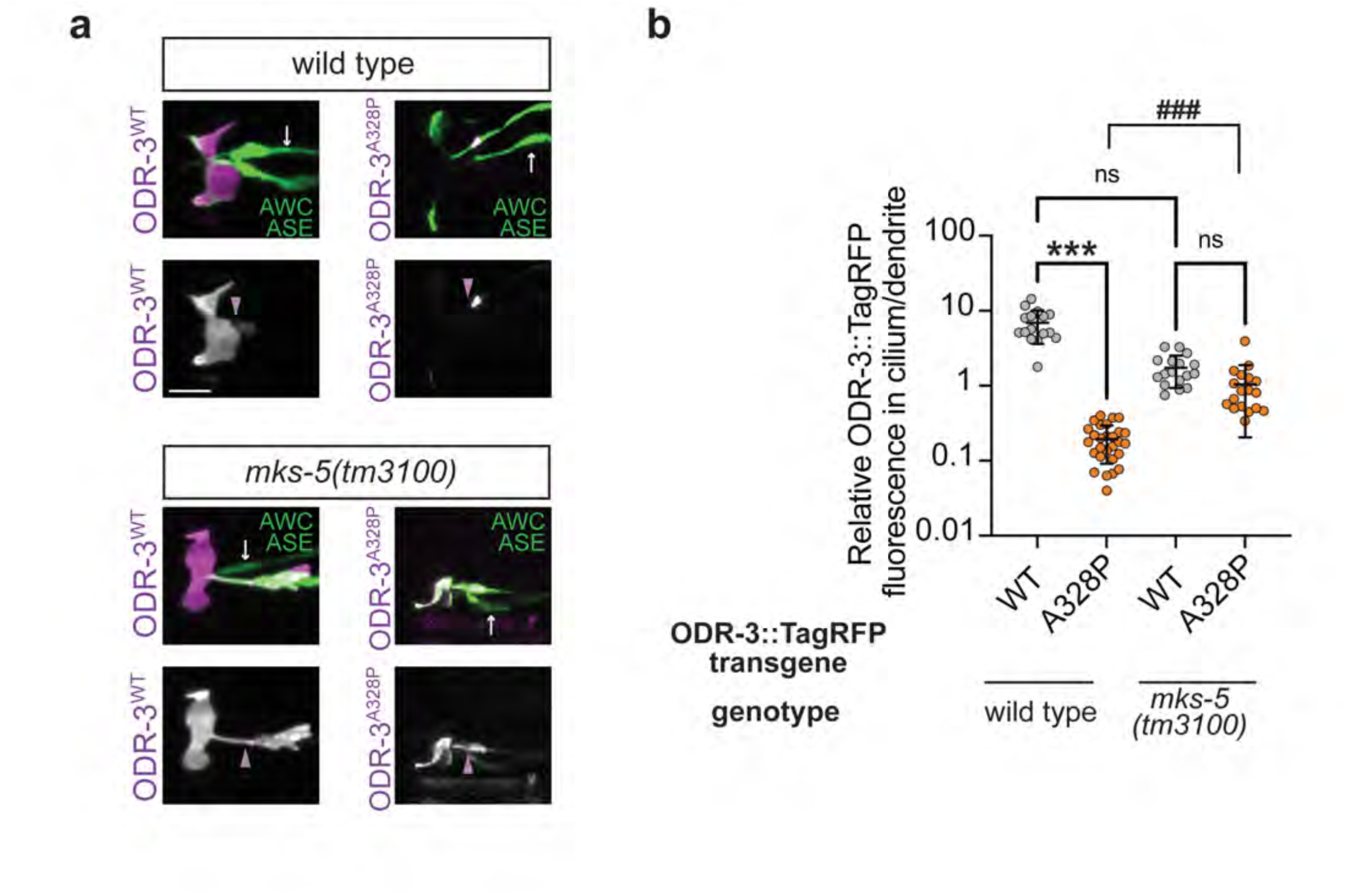
Intraciliary localization of A328P ODR-3 variant is increased in *mks-5* mutants. a) Representative images showing localization of TagRFP-tagged ODR-3^WT^ and ODR-3^A328P^ proteins in AWC cilia of wild-type animals (*top*) and *mks-5(tm3100)* mutants (*bottom*). AWC neurons were visualized via expression of *ceh-36*p::GFP, which also labels ASE neurons (white arrows). Purple arrowheads mark cilia base. Anterior is at left. Scale: 5 μm. b) Quantification of relative TagRFP fluorescence inside the AWC cilium vs distal dendrite for the indicated TagRFP-tagged ODR-3 variants in wild-type and *mks-5(tm3100)* mutant animals. Total number of AWC neurons analyzed for each variant: wild type (n=16 ODR-3^WT^, n=28 ODR-3^A328P^), *mks-5(tm3100)* (n=17 ODR-3^WT^, n=18 ODR-3^A328P^). Data for ODR-3^WT^ localization in the wild-type background are repeated from Fig. 3b. Means ± SD are indicated by horizontal and vertical bars, respectively. *** and ^###^ indicate different between bracketed groups at p<0.001; n.s.: not significant (Kruskal-Wallis with Dunn’s multiple comparisons test).

### *odr-3* mutations orthologous to patient *GNAI1* variants differentially affect AWC cilia morphology

To gain insight into the functional effects of the *GNAI1* disorder-associated variants *in vivo*, we used CRISPR-Cas9 editing to engineer worms homozygous for orthologous variants in the *odr-3* gene (Supplementary Fig. 4 and Supplementary Fig. 5). To visualize AWC cilia morphology, which is dependent on intact *odr-3* function, we then crossed an AWC-specific *gfp* reporter into the CRISPR-Cas9-engineered *odr-3* mutant strains. Consistent with published work, *odr-3(lf)* animals exhibited marked reduction in AWC cilia size compared to wild type (Fig. 5a and b). Cilia appeared normal in *M88V* and *I321T* homozygous animals (Fig. 5a and b). On the other hand, *T48I*, *D175V*, *A328P*, and *V334E* mutants exhibited cilia defects that ranged from mild albeit significant reduction in cilia size in *T48I* and *V334E* mutants to markedly smaller cilia in *D175V* and *A328P* mutants (Fig. 5a and b). Notably, the cilia phenotype in *A328P* animals was qualitatively and quantitatively indistinguishable from that of *odr-3(lf)* mutants, suggesting that *A328P* allele may be a functional null. Re-expression of wild-type *odr-3* in AWC neurons of *A328P* mutant animals partially but significantly rescued AWC cilia morphology (Fig. 5a and b), indicating that cilia defects in *A328P* mutants are indeed due to the loss of *odr-3* function. Despite multiple attempts, we were unable to engineer a *K272R* variant in the *odr-3* gene; so we took an alternative approach to determine whether this mutation impacts *odr-3* function. Specifically, we overexpressed ODR-3^K272R^ cDNA in AWC neurons of *odr-3(lf)* animals and quantified AWC cilia size in the transgenic animals. We found that these transgenic worms were indistinguishable from *odr-3(lf)* mutants (Fig. 5a and b), suggesting that *K272R* mutation strongly impairs ODR-3 activity.

**Fig. 5.**
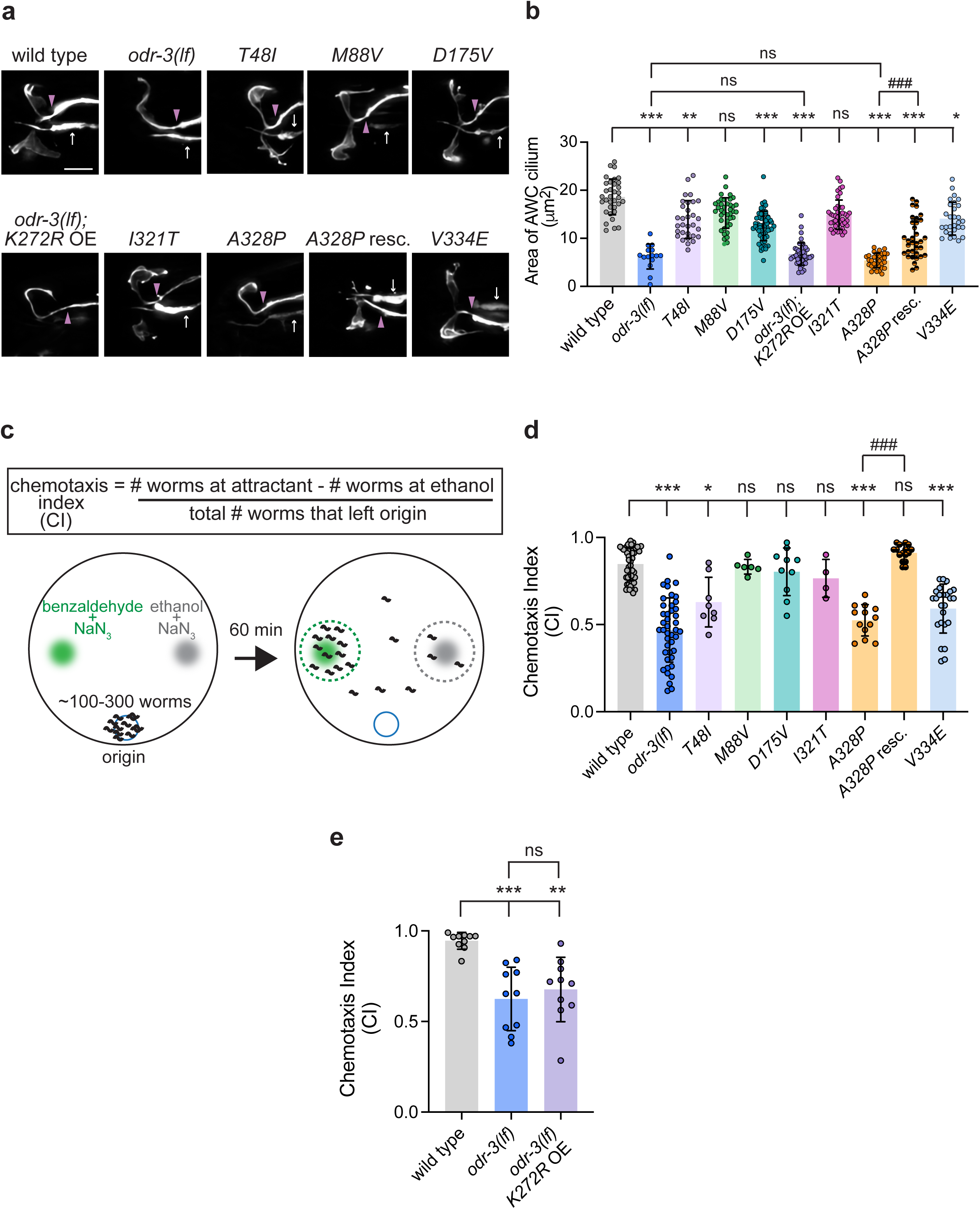
Orthologous patient mutations in the *odr-3* gene differentially alter AWC cilia morphology and function. a) Representative images of AWC cilia in animals homozygous for the indicated *odr-3* mutations and *odr-3(lf)* animals overexpressing the *odr-3^K272R^*transgene. AWC neurons were visualized via expression of *ceh-36*p::GFP, which also labels ASE neurons (white arrows). Purple arrowheads mark cilia base. Anterior is at left. Scale: 5 μm. b) Quantification of AWC cilia size in the indicated genotypes. Total number of cilia: WT (n=34), *odr-3(lf)* (n=15), *T48I* (n=31), *M88V* (n=40), *D175V* (n=59), *K272R* (n=45), *I321T* (n=41), *A328P* (n=32), *A328P* rescue (n=32), and *V334E* (n=29). Means ± SD are indicated by shaded and vertical bars, respectively. *, **, and *** indicate different from WT at p<0.05, 0.01, and 0.001, respectively (Kruskal-Wallis with Dunn’s multiple comparisons test). ^###^ indicates different between bracketed genotypes at p<0.001 (unpaired t test with Welch’s correction); n.s.: not significant (Mann Whitney test). c) Schematic of chemotaxis assay. d) Chemotaxis responses of the indicated homozygous variants. Individual data points represent chemotaxis index from one assay. Total number of assays: WT (n=50), *odr-3(lf)* (n=43), *T48I* (n=8), *M88V* (n=6), *D175V* (n=10), *I321T* (n=4), *A328P* (n=13), *A328P* rescue (n=16), and *V334E* (n=25). Means ± SD are indicated by shaded and vertical bars, respectively. * and *** indicate different from wild type at p<0.05 and 0.001, respectively; n.s.: not significant (Kruskal-Wallis with Dunn’s multiple comparisons test). ^###^ indicates different between bracketed genotypes at p<0.001 (unpaired t test with Welch’s correction). e) Chemotaxis responses of wild-type, *odr-3(lf)*, and *odr-3(lf)* mutants overexpressing *odr-3^K272R^* transgene in AWC neurons. Individual data points represent chemotaxis index from one assay. Total number of assays: WT (n=10), *odr-3(lf)* (n=10), and *odr-3(lf)*; *K272* OE (n=10). Means ± SD are indicated by shaded and vertical bars, respectively. ** and *** indicate different from wild type at p<0.01 and 0.001, respectively; n.s.: not significant (Kruskal-Wallis with Dunn’s multiple comparisons test).

*K272R* and *A328P* – two mutations that result in most severe cilia defects – alter conserved amino acids that come into direct contact with GDP/GTP. Lysine 272 participates in guanosine recognition (Luo et al., 2022), while alanine 328 is found in the TCAT motif, which is conserved throughout the G protein family, and forms a hydrogen bond with the oxygen in guanine (Luo et al., 2022). Therefore, cilia morphology defects observed in *K272R* and *A328P* mutants likely stem from aberrant interaction of ODR-3 with guanosine nucleotides.

### The *A328P* variant does not abolish ODR-3 interaction with RIC-8 or UNC-119

We previously showed that ODR-3 function is regulated by the cytoplasmic GEF and chaperone RIC-8 (Campagna et al., 2023). Like *odr-3(lf)* and *A328P* mutants, *ric-8(lf)* mutants exhibit a marked reduction in AWC cilia size. Since the *A328P* variant alters a residue just proximal to the Gα α5 helix, which participates in Gα – RIC-8 binding (Kant et al., 2016; Zeng et al., 2019), we next wanted to test whether this variant may impact cilia morphology by interfering with the ODR-3 – RIC-8 interaction. To address this question, we first examined *in vivo* association between ODR-3^A328P^ and RIC-8 via bimolecular fluorescence complementation (BiFC) (Shyu et al., 2008) and found that both ODR-3^WT^ and ODR-3^A328P^ variants tagged with the N-terminal Venus fragment (VN) associated with RIC-8 that was tagged with the C-terminal half of Venus (VC) (Supplementary Fig. 6a and b). Since BiFC is irreversible and may increase the likelihood of false positives over time (Kerppola, 2008; Xing et al., 2016), we next turned to FRET-FLIM (Algar et al., 2019) to confirm that ODR-3^A328P^ does indeed bind RIC-8 *in vivo*. We expressed GFP-fused RIC-8 alone or together with TagRFP-tagged ODR-3^WT^ or ODR-3^A328P^ constructs in AWC neurons under the ceh-36Δ promoter. As compared to RIC-8::GFP alone, the lifetime of GFP decreased in animals co-expressing RIC-8::GFP with ODR-3^WT^ confirming interaction between the two proteins (Supplementary Fig. 6c and d). Comparable reduction in GFP lifetime was also observed when we co-expressed RIC-8::GFP with ODR-3^A328P^ (Supplementary Fig. 6c and d), suggesting that *A328P* substitution does not abolish the association of the mutant ODR-3 with RIC-8, consistent with our BiFC data.

The GTP/GDP exchange cycle was also proposed to regulate the interaction between Gα proteins and an evolutionarily conserved trafficking adaptor UNC-119, which plays a key role in transporting Gα proteins to cilia (Zhang et al., 2011). Studies in mice and *C. elegans* demonstrated that *unc-119* deletion causes Gα protein mislocalization in sensory neurons of both species (Zhang et al., 2011). Furthermore, knocking down or mutating *unc-119* in zebrafish and *C. elegans*, respectively, produced phenotypes consistent with aberrant cilia function (Wright et al., 2011). So next we wondered whether *A328P* variant may disrupt ODR-3 interaction with UNC-119, as loss of this association could explain accumulation of ODR-3^A328P^ protein in the dendrite as well as cilia defects observed in *A328P* mutants. We expressed UNC-119 tagged with the C-terminal fragment of Venus (VC::UNC-119) together with either ODR-3^WT^ or ODR-3^A328P^ tagged with the N-terminal Venus fragment (ODR-3^WT^::VN and ODR-3^A328P^::VN, respectively) in AWC neurons of wild-type animals. We observed BiFC in 28% of animals co-expressing VC::UNC-119 and free N-terminal Venus fragment (negative control) (Supplementary Fig. 7). Co-expression of VC::UNC-119 with ODR-3^WT^::VN or ODR-3^A328P^::VN reconstituted fluorescent signal in AWC neurons of a significantly greater percent of transgenic animals (81% and 51%, respectively) (Supplementary Fig. 7b), indicating that ODR-3^A328P^ retains ability to interact with UNC-119. However, the BiFC signal in animals co-expressing VC::UNC-119 with ODR-3^A328P^::VN appeared to concentrate at the cilia base and was qualitatively weaker inside the cilium of AWC neurons compared to animals co-expressing VC::UNC-119 and ODR-3^WT^::VN (Supplementary Fig. 7a). These results are consistent with the hypothesis that ciliary transport of the ODR-3-UNC-119 complex may be compromised by the *A328P* mutation – a finding that is consistent with accumulation of the endogenously tagged ODR-3^A328P^::split-wrmScarlet at cilia base (see Supplementary Fig. 3).

### The *T48I*, *K272R*, and *A328P* variants in the guanine-binding domains and *V334E* variant in the putative receptor-binding motif impair AWC-mediated chemotaxis

We next examined the impact of orthologous variants on sensory responses mediated by AWC. Ciliated endings of AWC neurons are enveloped in the glial processes (Doroquez et al., 2014; Perkins et al., 1986) and house signaling machinery that detects a panel of volatile attractants including benzaldehyde (Bargmann et al., 1993; Ferkey et al., 2021). However, prior studies reported that these primary behavioral responses to volatile odorants are preserved in several mutants with severely compromised cilia morphology (Campagna et al., 2023; Philbrook et al., 2024), thereby uncoupling cilia morphology from cilia function in mediating odorant sensing by AWC. To investigate the impact of orthologous patient variants on ODR-3-dependent attraction toward benzaldehyde, we carried out population chemotaxis assays (Fig. 5c). As expected, wild-type animals exhibited robust attraction to benzaldehyde (1:200), while *odr-3(lf)* animals were deficient in this response (Roayaie et al., 1998) (Fig. 5d). *M88V* and *I321T* mutants exhibited normal chemotaxis, suggesting that these variants do not impair ODR-3 function in either AWC ciliogenesis or chemotaxis toward benzaldehyde. On the other hand, *D175V* mutants exhibited normal behavioral response (Fig. 5d), despite having morphologically defective AWC cilia (see Fig. 5a and b). Normal chemotaxis of *D175V* mutants was particularly intriguing given the dramatic impact of this variant on ciliary localization of the encoded ODR-3 protein (see Fig. 3a) and suggests that ODR-3 enrichment inside the AWC cilium may be important for its role in mediating ciliogenesis but not benzaldehyde attraction.

The variants mapping to the nucleotide-binding domain all resulted in impaired chemotaxis toward benzaldehyde. Specifically, *A328P* and *T48I* homozygous mutants as well as transgenic animals expressing *odr-3^K272R^* cDNA in *odr-3(lf)* AWC neurons exhibited significantly lower chemotaxis indices compared to wild type (Fig. 5d and e), consistent with the hypothesis that these variants disrupt overall ODR-3 function. Of note, the *V334E* variant, which alters an amino acid in the putative GPCR-interaction motif, also negatively impacted ODR-3-mediated behavior (Fig. 5d), suggesting that this missense mutation may disrupt ODR-3 association with GPCRs. The summary of all cellular and behavioral phenotypes exhibited by the orthologous patient variants is provided in Table 1.

**Table 1:**
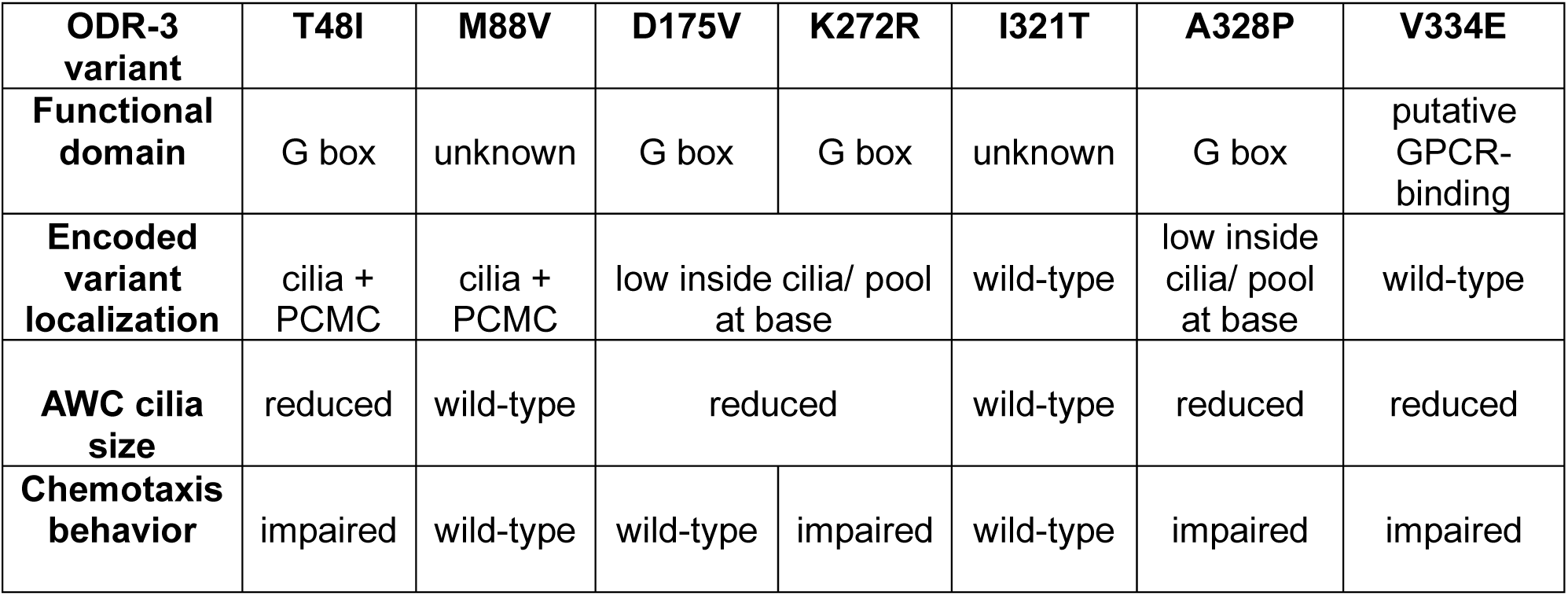
Summary of AWC cellular and behavioral phenotypes caused by orthologous mutations in the *odr-3* gene.

| **ODR-3 variant** | **T48I** | **M88V** | **D175V** | **K272R** | **I321T** | **A328P** | **V334E** |
| --- | --- | --- | --- | --- | --- | --- | --- |
| **Functional domain** | G box | unknown | G box | G box | unknown | G box | putative GPCR-binding |
| **Encoded variant localization** | cilia + PCMC | cilia + PCMC | low inside cilia/ pool at base |  | wild-type | low inside cilia/ pool at base | wild-type |
| **AWC cilia size** | reduced | wild-type | reduced |  | wild-type | reduced | reduced |
| **Chemotaxis behavior** | impaired | wild-type | wild-type | impaired | wild-type | impaired | impaired |

### The *D175V* variant disrupts ODR-3 ciliary localization and sensory function of ASH neurons

In addition to AWC, *odr-3* is expressed in several sensory neurons including the nociceptor ASH that detects a range of aversive chemicals (Roayaie et al., 1998). Similarly to AWC, *odr-3* mediates chemosensory transduction in ASH neurons (Bargmann et al., 1993; Roayaie et al., 1998; Yoshida et al., 2012). However, *odr-3* function appears to be dispensable for ASH cilium assembly (Roayaie et al., 1998) (Supplementary Fig. 8), pointing to cell-specific roles for this gene in cilia biology.

Unlike AWC, ASH neurons possess simple rod-like cilia that extend through a glial channel and are exposed to the external environment (Doroquez et al., 2014; Perkins et al., 1986) (Fig. 6a). A recent study demonstrated that a minimum ASH cilium length is required for ASH-mediated sensory responses to aqueous chemicals, including behavioral avoidance of high-osmolarity solutions (Philbrook et al., 2024). In this context, the minimum length is likely necessary to allow for direct contact between aqueous cues and cilia-localized signaling machinery. Therefore, we wondered whether the *D175V* variant, which excludes the encoded ODR-3 protein from the cilium but does not impair chemotaxis in AWC neurons, may have a different effect on the ASH cilium function. First, we confirmed that TagRFP-tagged ODR-3^WT^ expressed under the *sra-6* regulatory sequences localized to ASH cilia in wild-type animals (Fig. 6b). Like in AWC, ODR-3^D175V^ expressed in ASH neurons of wild type *C. elegans* was largely excluded from cilia and instead accumulated in the distal dendrite (Fig. 6b and c), suggesting that *D175V* mutation interferes with ODR-3 trafficking mechanisms conserved across cell types.

**Fig. 6.**
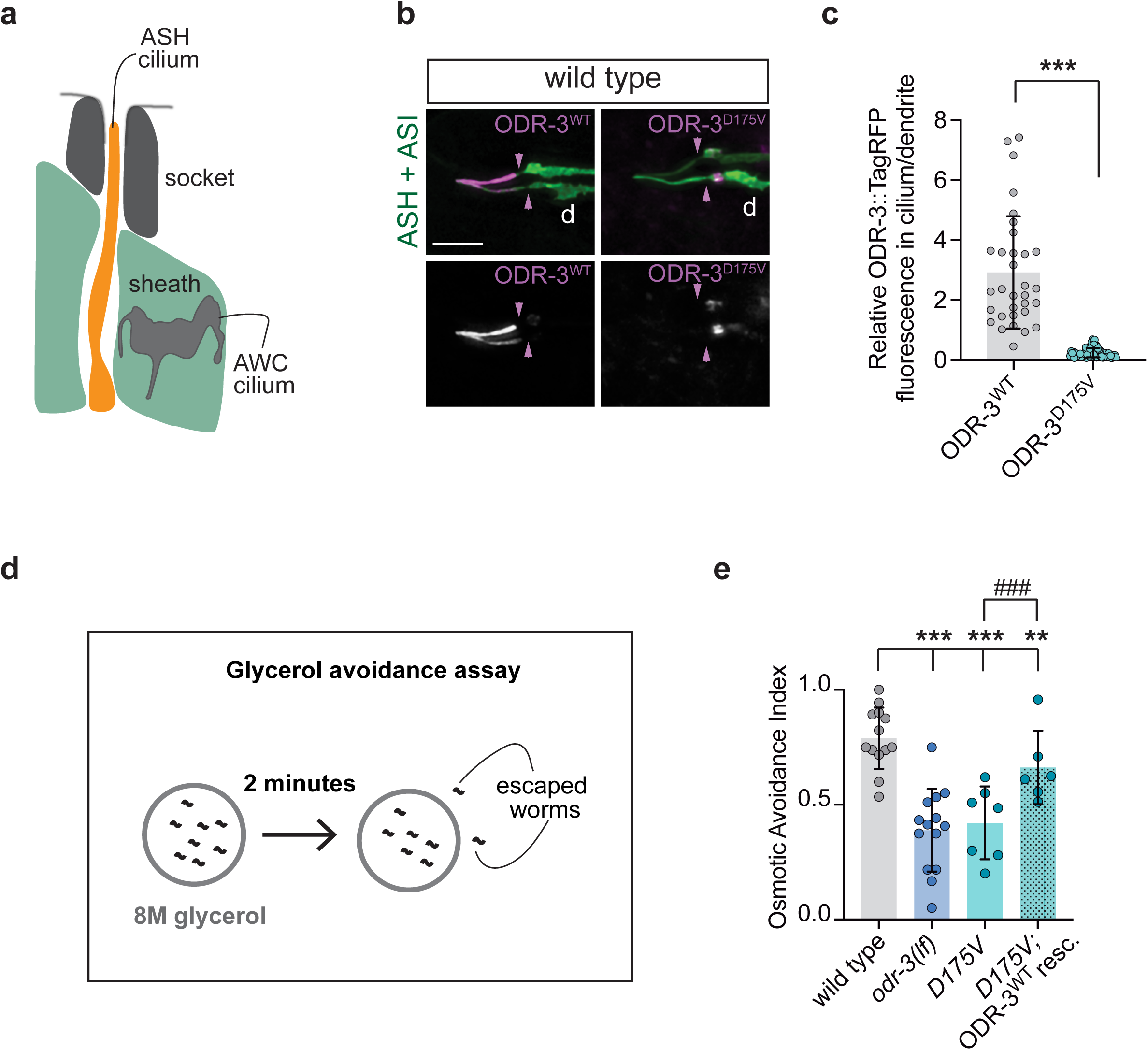
The *D175V* variant disrupts ODR-3 localization to the ASH cilia and impairs ASH-mediated behavior. a) Schematic of the amphid channel. Only AWC and ASH cilia are shown for simplicity. Socket and sheath are glial cells that form the channel. Anterior is up. b) Representative images showing localization of TagRFP-tagged WT and D175V variants of ODR-3 in ASH and ASI neurons. ASH and ASI neurons were visualized via expression of *sra-6*p::GFP. Purple arrowheads mark cilia base. Anterior is at left. Scale: 5 μm. c) Quantification of relative TagRFP fluorescence inside the ASH cilium vs distal dendrite for the indicated TagRFP-tagged ODR-3 variants. Total number of ASH neurons analyzed for each variant: WT (n=32), *D175V* (n=65). *** indicates different from WT at p<0.001 (Mann-Whitney test). d) Diagram of glycerol avoidance assay. e) Fraction of animals that remain inside of an 8M-glycerol ring after two minutes (osmotic avoidance index). Genotypes of assayed animals are indicated. Each data point represents a mean osmotic index per genotype per day. Three to eight technical replicates were assayed per genotype per day. Total number of assayed animals: WT (n=298), *odr-3(lf)* (n=264), *D175V* (n=224), *D175* rescue (n=157). Means ± SD are indicated by shaded and vertical bars, respectively. ** and *** indicate different from wild type at p<0.01 and 0.001, respectively (Fisher’s exact test). ^###^ indicates different between bracketed genotypes (Fisher’s exact test).

Next, we examined whether ODR-3 localization to the ASH cilium is required for its function in mediating avoidance of high-osmolarity solutions. To this end, we placed wild-type, *odr-3(lf)*, or *D175V* homozygous animals inside a ring of 8M glycerol and counted the number of animals that escaped the glycerol ring within two minutes (Cornils et al., 2016) (Fig. 6d). On average, 80% of wild-type animals avoided glycerol and stayed inside the ring (Fig. 6e). In contrast, *odr-3(lf)* mutants escaped the ring, and only 40% of *odr-3(lf)* animals remained inside the ring after two minutes (Fig. 6e). Similarly to *odr-3(lf)* animals, 42% of *D175V* mutants remained inside the glycerol ring after 2 minutes (Fig. 6e), indicating that *D175V* variant impairs ASH sensory function. Importantly, over-expression of wild-type *odr-3* cDNA in ASH neurons of *D175V* animals partially but significantly rescued glycerol avoidance (Fig. 6e), confirming that the observed sensory deficit is not due to a second-site mutation.

### D173V, K270R, and A326P patient variants impair ciliary localization of Gαi1 protein in human cells

We next explored whether the variants with the strongest impact on ODR-3 localization demonstrated similar localization defects in human cells. Gαi1 was previously reported to localize to primary cilia of cultured mouse fibroblasts (Singh et al., 2015). We first examined sub-cellular localization of wild-type human Gαi1 by transfecting HEK293T cells with *eGFP-* tagged *GNAI1* cDNA. In line with the published data, human Gαi1^WT^::eGFP was present inside the cilia as well as at the plasma membrane and Golgi (Gabay et al., 2011) (Fig. 7a and b). Strikingly, *D173V*, *K270R*, and *A326P* patient variants orthologous to the examined ODR-3 mutants significantly reduced intra-ciliary levels of the encoded proteins relative to Golgi (Fig. 7a and b), demonstrating that the impact of these variants on Gα protein trafficking is evolutionarily conserved.

**Fig. 7.**
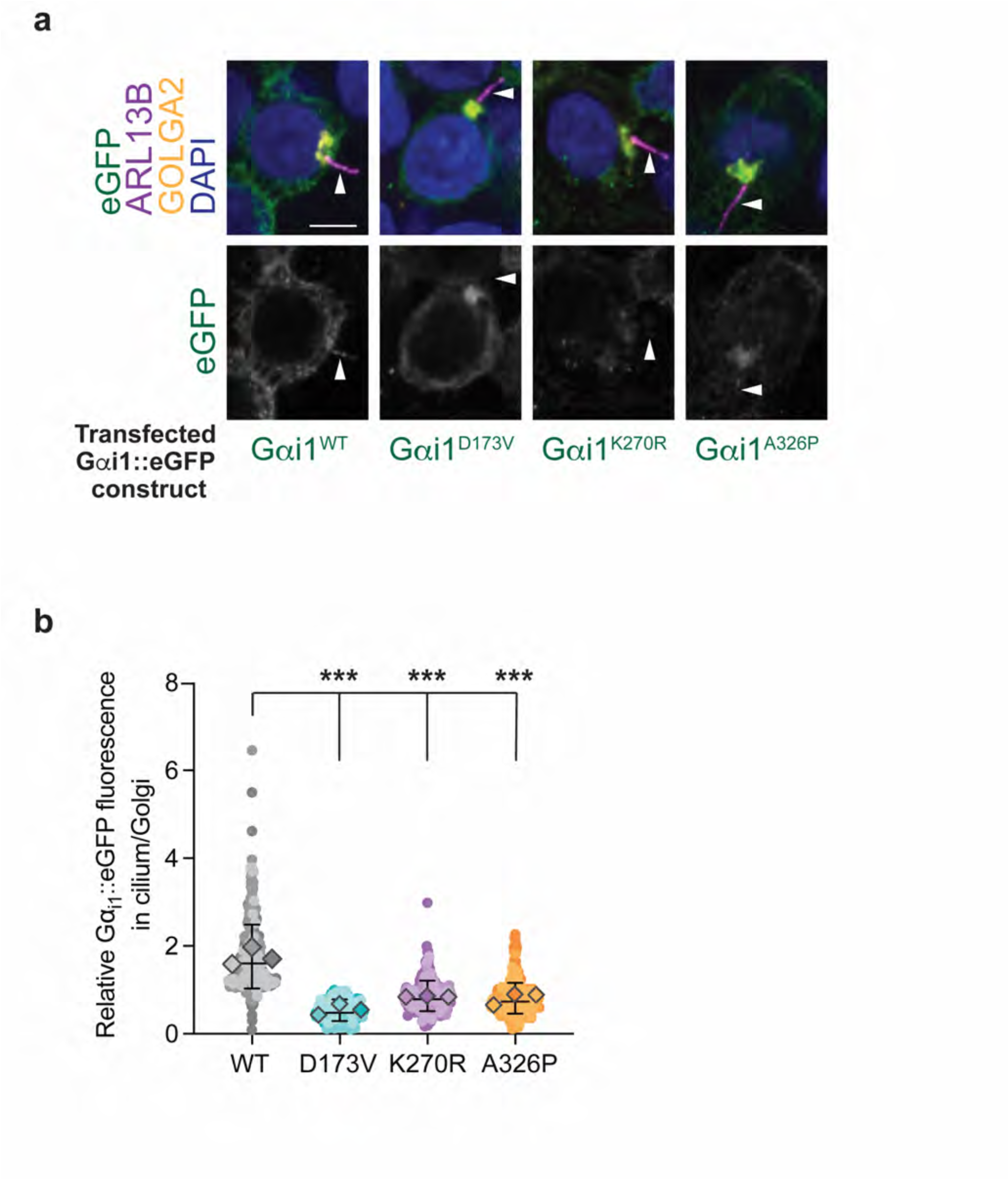
The *D173V*, *K270R*, and *A326P* variants in *GNAI1* impair cilia localization of the encoded proteins in human cells. a) Immunofluorescence images of fixed HEK293T cells transfected with the indicated eGFP-tagged Gαi1 variants. White arrowheads mark cilia labeled with anti-ARL13B antibody. DAPI and anti-GOLGA2 label DNA and Golgi, respectively. Scale: 5 μm. b) Quantification of relative eGFP fluorescence in cilium vs Golgi for the indicated eGFP-tagged Gαi1 variants. Individual biological replicates for each variant are shown in different shades of the corresponding color. Diamonds: mean of individual biological replicates. Total mean ± SD across all biological replicates is indicated by horizontal and vertical bars, respectively. Total number of cells analyzed for each Gαi1 variant: WT (n=322), *D173V* (n=220), *K270R* (n=260), *A326P* (n=315). *** indicates different from Gαi1^WT^ at p<0.001 (Kruskal-Wallis with Dunn’s multiple comparisons test).

## DISCUSSION

### Orthologous *GNAI1*-disorder patient variants differentially impact *odr-3* function

Here, we demonstrate that *GNAI1* – the causal gene in a recently identified neurodevelopmental disorder – regulates ciliogenesis in two human cell lines, suggesting that cilia dysfunction may play a role in the pathogenesis of *GNAI1* disorder. We used *C. elegans* as a whole-animal model to functionally classify a subset of orthologous *GNAI1* disorder mutations *in vivo* and found variant- and cell-specific effects on the examined phenotypes. Overall, orthologous missense mutations that altered conserved amino-acid residues in the nucleotide-binding interface of Gα (ODR-3: *T48I*, *K272R*, and *A328P*) caused defects in both examined ODR-3-dependent phenotypes – AWC cilia morphology and chemotaxis behavior. Similarly, the *V334E* variant, which maps to the terminal α5 helix of Gα ODR-3, caused a mild reduction in AWC cilia size and significantly impaired attraction to benzaldehyde. The α5 helix of Gα proteins participates in GPCR binding and receptor-catalyzed nucleotide exchange (Alexander et al., 2014; Marin et al., 2002; Masuho et al., 2023; Oldham et al., 2006). Furthermore, the *V332A* substitution was shown to significantly destabilized the GDP-bound state of Gαi1 (Sun et al., 2015). Therefore, additional *in vitro* studies could shed more light on GPCR binding efficiency and nucleotide exchange rates of *V332E* Gαi1 variant present in *GNAI1* disorder patients.

On the other hand, variants altering conserved amino acids in domains of unknown function (ODR-3: *M88V* and *I321T*) had no impact on either AWC cilia morphology or behavior. Recent studies have begun to highlight challenges associated with interpreting pathogenicity of *de novo* variants across developmental disorders (Mani, 2017). For example, one study estimated that only ∼13% of *de novo* missense variants contribute to risk of autism-spectrum disorder (Iossifov et al., 2014). Therefore, functional studies in additional *in vivo* models are needed to confirm the potentially benign or disease-contributing nature of *M88V* and *I321T* variants.

Intriguingly, the *D175V* variant, which caused marked mislocalization of the encoded ODR-3 protein from the cilium to the distal dendrite, was associated with significantly reduced AWC cilia size, yet had no impact on AWC-mediated attraction to benzaldehyde. In stark contrast, the same *D175V* variant did not affect cilia length in ASH neurons but compromised ASH-mediated glycerol avoidance, demonstrating that the functional impact of the same variant varies depending on the cellular context. Context-dependent differences in functional effects of patient variants have also been reported for other disorders (Di Rocco et al., 2023; Feng et al., 2017; Muntean et al., 2021; Wang et al., 2022; Wong et al., 2019), highlighting the importance of modeling patient mutations in multiple experimental systems and disease-relevant contexts to gain a comprehensive understanding of the underlying disease mechanisms.

### Modeling orthologous *GNAI1*-disorder missense variants in *C. elegans* provides additional insight into cell-specific functions of ODR-3

Three of the examined orthologous variants (ODR-3: *D175V*, *K272R*, and *A328P*) caused qualitatively and quantitatively similar defects in localization of the encoded ODR-3 proteins. Specifically, mutant ODR-3 variants accumulated in the distal dendrite and were present at markedly reduced levels inside the AWC cilium, where wild-type ODR-3 normally resides. Both *K272R* and *A328P* variants alter amino acids that come into direct contact with the nucleotide. Although the *D175* residue does not directly bind GDP, it is found in the immediate vicinity of the nucleotide-binding motif (see Fig. 2) (Kant et al., 2016). Prior *in vitro* work showed that a different missense mutation at the A326 position of Gαi1 (*A326S*) accelerated dissociation of GDP from the αi1β1γ1 heterotrimer by > 200-fold (Posner et al., 1998), suggesting that proper confirmation of the nucleotide-binding pocket or GDP binding itself may be necessary for efficient translocation of ODR-3/Gαi1 into the cilium. Improved targeting of ODR-3^A328P^ protein to the cilia in *mks-5(lf)* mutants is consistent with the hypothesis that *A328P* mutation impedes translocation of ODR-3 across the TZ. However, based on the results from the BiFC assay, ODR-3^A328P^ appears to still interact with UNC-119 – a key mediator of Gα ciliary transport – suggesting that additional mechanisms may contribute to Gα trafficking into the cilium. Since the BiFC signal is binary (either on or off), in the future, it would be worthwhile to directly examine the stability of UNC-119 complexes with ODR-3/Gαi1 proteins carrying *D175V/D173V*, *K272R/K270R*, and *A328P/A326P* patient substitutions using biochemical approaches.

We also found that although the *D175V* variant caused mislocalization of the encoded ODR-3 protein from the cilium to the dendrite, it did not alter chemotaxis toward benzaldehyde. This observation led us to conclude that ODR-3 localization to the cilium is not necessary for its function in mediating primary behavioral responses to volatile odorants detected by AWC. Similarly, a recent study demonstrated that diacetyl receptor ODR-10 and Gα ODR-3, which also functions downstream from ODR-10 to mediate attraction toward diacetyl, translocate from the cilium to dendritic branches in AWA olfactory neurons upon disruption of intraflagellar transport (IFT) (Philbrook et al., 2024). Interestingly, IFT mutants retain normal primary responses to diacetyl but exhibit defects in habituation and desensitization (Larsch et al., 2015; Philbrook et al., 2024). It will be of interest to determine whether sequestration of ODR-3^D175V^ in the distal dendrite similarly impacts desensitization and habituation responses of AWA and AWC sensory neurons.

Finally, in contrast to AWC, ODR-3 localization to the ASH cilium is required for its function in glycerol avoidance, as *D175V* homozygous mutants exhibit marked deficits in this ASH-mediated sensory behavior despite having cilia of normal length. Recent electron microscopy studies in the human and mouse cortex uncovered remarkable complexity of cilia interactions with the cellular environment in the brain (Ott et al., 2024; Wu et al., 2024). Neuronal cilia were shown to be immersed into dense neuropil and adjacent to axonal and dendritic processes at several points along their length (Ott et al., 2024). Occasionally, tips of neuronal cilia were also found to be enveloped by astrocytes. While the exact nature and functional significance of these interactions remain a mystery, synapses were frequently observed adjacent to cilia in mouse and human cortical tissue (Ott et al., 2024; Wu et al., 2024), and one study even reported axo-ciliary synapses between brainstem axons and cilia of hippocampal CA1 neurons (Sheu et al., 2022). Collectively, these observations propose an exciting hypothesis that primary cilia may be a major player in neural circuit modulation. Therefore, mislocalization of Gαi1 – a key transducer of GPCR signaling that modulates many aspects of neuronal physiology – from cilia to ectopic sub-cellular locations may have a sizeable impact on local neural connectivity as well as neuron-neuron and/or neuron-glia communication. It would be worthwhile to determine whether the patient Gαi1 variants that exhibit reduced ciliary levels in *C. elegans* neurons and human epithelial cells are similarly excluded from cilia of distinct neuron classes in the mammalian brain and to classify their impact on neuronal cilia length and circuit function.

In conclusion, our results highlight the power of *C. elegans* as a whole-organism system to rapidly determine functional impact of missense patient variants in conserved Gα residues and thus prioritize functionally consequential mutations for further mechanistic studies in mammalian models and/or additional disease-relevant contexts. Moreover, orthologous disease-associated mutations in model organisms like *C. elegans* can provide new insights into normal function of the impacted genes.

## Data availability

Strains and plasmids used in this work are listed in Supplementary Tables 1 and 2, respectively, and are available upon request. All data necessary for confirming the conclusions of the article are present within the article, figures, table, and supplementary files.

## Acknowledgements

We are grateful to members of the Nechipurenko lab for critical comments on the manuscript and to Ryan Breitenbach for technical assistance and maintenance of strains. Some strains were provided by the CGC, which is funded by NIH Office of Research Infrastructure Programs (P40 OD010440).

## REFERENCES

Alexander, N. S., Preininger, A. M., Kaya, A. I., Stein, R. A., Hamm, H. E., & Meiler, J. (2014). Energetic analysis of the rhodopsin-G-protein complex links the alpha5 helix to GDP release. Nat Struct Mol Biol, 21(1), 56–63. 10.1038/nsmb.2705

Algar, W. R., Hildebrandt, N., Vogel, S. S., & Medintz, I. L. (2019). FRET as a biomolecular research tool — understanding its potential while avoiding pitfalls. Nature Methods, 16(9), 815–829. 10.1038/s41592-019-0530-8

Anvarian, Z., Mykytyn, K., Mukhopadhyay, S., Pedersen, L. B., & Christensen, S. T. (2019). Cellular signalling by primary cilia in development, organ function and disease. Nature Reviews Nephrology, 15(4), 199–219. 10.1038/s41581-019-0116-9

Bargmann, C. I., Hartwieg, E., & Horvitz, H. R. (1993). Odorant-selective genes and neurons mediate olfaction in C. elegans. Cell, 74(3), 515–527. 10.1016/0092-8674(93)80053-h

Betke, K. M., Wells, C. A., & Hamm, H. E. (2012). GPCR mediated regulation of synaptic transmission. Prog Neurobiol, 96(3), 304–321. 10.1016/j.pneurobio.2012.01.009

Brenner, S. (1974). The genetics of Caenorhabditis elegans. Genetics, 77(1), 71–94. 10.1093/genetics/77.1.71

Campagna, C. M., McMahon, H., & Nechipurenko, I. (2023). The G protein alpha chaperone and guanine-nucleotide exchange factor RIC-8 regulates cilia morphogenesis in Caenorhabditis elegans sensory neurons. PLoS Genetics, 19(11), e1011015. 10.1371/journal.pgen.1011015

Cevik, S., Sanders, A. A., Van Wijk, E., Boldt, K., Clarke, L., van Reeuwijk, J., Hori, Y., Horn, N., Hetterschijt, L., Wdowicz, A., Mullins, A., Kida, K., Kaplan, O. I., van Beersum, S. E., Man Wu, K., Letteboer, S. J., Mans, D. A., Katada, T., Kontani, K., . . . Blacque, O. E. (2013). Active transport and diffusion barriers restrict Joubert Syndrome-associated ARL13B/ARL-13 to an Inv-like ciliary membrane subdomain. PLoS Genet, 9(12), e1003977. 10.1371/journal.pgen.1003977

Cornils, A., Maurya, A. K., Tereshko, L., Kennedy, J., Brear, A. G., Prahlad, V., Blacque, O. E., & Sengupta, P. (2016). Structural and Functional Recovery of Sensory Cilia in C. elegans IFT Mutants upon Aging. PLoS Genetics, 12(12), e1006325. 10.1371/journal.pgen.1006325

Culotti, J. G., & Russell, R. L. (1978). Osmotic avoidance defective mutants of the nematode Caenorhabditis elegans. Genetics, 90(2), 243–256. 10.1093/genetics/90.2.243

Di Rocco, M., Galosi, S., Follo, F. C., Lanza, E., Folli, V., Martire, A., Leuzzi, V., & Martinelli, S. (2023). Phenotypic Assessment of Pathogenic Variants in GNAO1 and Response to Caffeine in C. elegans Models of the Disease. Genes (Basel), 14(2). 10.3390/genes14020319

Dokshin, G. A., Ghanta, K. S., Piscopo, K. M., & Mello, C. C. (2018). Robust Genome Editing with Short Single-Stranded and Long, Partially Single-Stranded DNA Donors in Caenorhabditis elegans. Genetics, 210(3), 781–787. 10.1534/genetics.118.301532

Doroquez, D. B., Berciu, C., Anderson, J. R., Sengupta, P., & Nicastro, D. (2014). A high-resolution morphological and ultrastructural map of anterior sensory cilia and glia in Caenorhabditis elegans. eLife, 3. 10.7554/elife.01948

Dror, R. O., Mildorf, T. J., Hilger, D., Manglik, A., Borhani, D. W., Arlow, D. H., Philippsen, A., Villanueva, N., Yang, Z., Lerch, M. T., Hubbell, W. L., Kobilka, B. K., Sunahara, R. K., & Shaw, D. E. (2015). SIGNAL TRANSDUCTION. Structural basis for nucleotide exchange in heterotrimeric G proteins. Science, 348(6241), 1361–1365. 10.1126/science.aaa5264

Feng, H., Sjögren, B., Karaj, B., Shaw, V., Gezer, A., & Neubig, R. R. (2017). Movement disorder in GNAO1 encephalopathy associated with gain-of-function mutations. Neurology, 89(8), 762–770. 10.1212/wnl.0000000000004262

Ferkey, D. M., Sengupta, P., & L’Etoile, N. D. (2021). Chemosensory signal transduction in Caenorhabditis elegans. Genetics, 217(3). 10.1093/genetics/iyab004

Gabay, M., Pinter, M. E., Wright, F. A., Chan, P., Murphy, A. J., Valenzuela, D. M., Yancopoulos, G. D., & Tall, G. G. (2011). Ric-8 Proteins Are Molecular Chaperones That Direct Nascent G Protein Subunit Membrane Association. Science Signaling, 4(200), ra79-ra79. 10.1126/scisignal.2002223

Garcia-Gonzalo, F. R., & Reiter, J. F. (2017). Open Sesame: How Transition Fibers and the Transition Zone Control Ciliary Composition. Cold Spring Harbor Perspectives in Biology, 9(2), a028134. 10.1101/cshperspect.a028134

Gilman, A. G. (1987). G PROTEINS: TRANSDUCERS OF RECEPTOR-GENERATED SIGNALS. Annual Review of Biochemistry, 56(1), 615–649. 10.1146/annurev.bi.56.070187.003151

Goudeau, J., Sharp, C. S., Paw, J., Savy, L., Leonetti, M. D., York, A. G., Updike, D. L., Kenyon, C., & Ingaramo, M. (2021). Split-wrmScarlet and split-sfGFP: tools for faster, easier fluorescent labeling of endogenous proteins in Caenorhabditis elegans. Genetics, 217(4). 10.1093/genetics/iyab014

Guemez-Gamboa, A., Coufal, N. G., & Gleeson, J. G. (2014). Primary cilia in the developing and mature brain. Neuron, 82(3), 511–521. 10.1016/j.neuron.2014.04.024

Hamada, N., Iwamoto, I., Kawamura, N., & Nagata, K. I. (2021). Heterotrimeric G-protein, Gi1, is involved in the regulation of proliferation, neuronal migration, and dendrite morphology during cortical development in vivo. J Neurochem, 157(4), 1167-1181. 10.1111/jnc.15205

Hasenpusch-Theil, K., & Theil, T. (2021). The Multifaceted Roles of Primary Cilia in the Development of the Cerebral Cortex. Front Cell Dev Biol, 9, 630161. 10.3389/fcell.2021.630161

Hildebrandt, F., Benzing, T., & Katsanis, N. (2011). Ciliopathies. N Engl J Med, 364(16), 1533–1543. 10.1056/NEJMra1010172

Hilgendorf, K. I., Johnson, C. T., & Jackson, P. K. (2016). The primary cilium as a cellular receiver: organizing ciliary GPCR signaling. Current Opinion in Cell Biology, 39, 84–92. 10.1016/j.ceb.2016.02.008

Iossifov, I., O’Roak, B. J., Sanders, S. J., Ronemus, M., Krumm, N., Levy, D., Stessman, H. A., Witherspoon, K. T., Vives, L., Patterson, K. E., Smith, J. D., Paeper, B., Nickerson, D. A., Dea, J., Dong, S., Gonzalez, L. E., Mandell, J. D., Mane, S. M., Murtha, M. T., . . . Wigler, M. (2014). The contribution of de novo coding mutations to autism spectrum disorder. Nature, 515(7526), 216–221. 10.1038/nature13908

Jensen, V. L., Li, C., Bowie, R. V., Clarke, L., Mohan, S., Blacque, O. E., & Leroux, M. R. (2015). Formation of the transition zone by Mks5/Rpgrip1L establishes a ciliary zone of exclusion (CIZE) that compartmentalises ciliary signalling proteins and controls PIP2 ciliary abundance. The EMBO Journal, 34(20), 2537–2556. 10.15252/embj.201488044

Jurisch-Yaksi, N., Wachten, D., & Gopalakrishnan, J. (2024). The neuronal cilium - a highly diverse and dynamic organelle involved in sensory detection and neuromodulation. Trends Neurosci, 47(5), 383–394. 10.1016/j.tins.2024.03.004

Kant, R., Zeng, B., Thomas, C. J., Bothner, B., & Sprang, S. R. (2016). Ric-8A, a G protein chaperone with nucleotide exchange activity induces long-range secondary structure changes in Gα. eLife, 5. 10.7554/elife.19238

Karalis, V., Donovan, K. E., & Sahin, M. (2022). Primary Cilia Dysfunction in Neurodevelopmental Disorders beyond Ciliopathies. J Dev Biol, 10(4). 10.3390/jdb10040054

Kerppola, T. K. (2008). Bimolecular Fluorescence Complementation (BiFC) Analysis as a Probe of Protein Interactions in Living Cells. Annual Review of Biophysics, 37(1), 465–487. 10.1146/annurev.biophys.37.032807.125842

Kim, K., Kim, R., & Sengupta, P. (2010). The HMX/NKX homeodomain protein MLS-2 specifies the identity of the AWC sensory neuron type via regulation of the ceh-36 Otx gene in C. elegans. Development, 137(6), 963–974. 10.1242/dev.044719

Kurabayashi, N., Nguyen, M. D., & Sanada, K. (2013). The G protein-coupled receptor GPRC5B contributes to neurogenesis in the developing mouse neocortex. Development, 140(21), 4335–4346. 10.1242/dev.099754

Lange, K. I., Best, S., Tsiropoulou, S., Berry, I., Johnson, C. A., & Blacque, O. E. (2022). Interpreting ciliopathy-associated missense variants of uncertain significance (VUS) in Caenorhabditis elegans. Hum Mol Genet, 31(10), 1574–1587. 10.1093/hmg/ddab344

Lange, K. I., Tsiropoulou, S., Kucharska, K., & Blacque, O. E. (2021). Interpreting the pathogenicity of Joubert syndrome missense variants in Caenorhabditis elegans. Dis Model Mech, 14(1). 10.1242/dmm.046631

Lans, H., Rademakers, S., & Jansen, G. (2004). A Network of Stimulatory and Inhibitory Gα-Subunits Regulates Olfaction in Caenorhabditis elegans. Genetics, 167(4), 1677–1687. 10.1534/genetics.103.024786

Larsch, J., Flavell, S. W., Liu, Q., Gordus, A., Albrecht, D. R., & Bargmann, C. I. (2015). A Circuit for Gradient Climbing in C. elegans Chemotaxis. Cell Rep, 12(11), 1748–1760. 10.1016/j.celrep.2015.08.032

Lee, J. E., & Gleeson, J. G. (2011). Cilia in the nervous system: linking cilia function and neurodevelopmental disorders. Curr Opin Neurol, 24(2), 98–105. 10.1097/WCO.0b013e3283444d05

Luo, M., Han, Z., Huang, G., Li, R., Liu, Y., Lu, J., Liu, L., & Miao, R. (2022). Structural comparison of unconventional G protein YchF with heterotrimeric G protein and small G protein. Plant Signal Behav, 17(1), 2024405. 10.1080/15592324.2021.2024405

Mani, A. (2017). Pathogenicity of De Novo Rare Variants: Challenges and Opportunities. Circ Cardiovasc Genet, 10(6). 10.1161/CIRCGENETICS.117.002013

Marin, E. P., Krishna, A. G., & Sakmar, T. P. (2002). Disruption of the alpha5 helix of transducin impairs rhodopsin-catalyzed nucleotide exchange. Biochemistry, 41(22), 6988–6994. 10.1021/bi025514k

Masuho, I., Kise, R., Gainza, P., Von Moo, E., Li, X., Tany, R., Wakasugi-Masuho, H., Correia, B. E., & Martemyanov, K. A. (2023). Rules and mechanisms governing G protein coupling selectivity of GPCRs. Cell Rep, 42(10), 113173. 10.1016/j.celrep.2023.113173

Muir, A. M., Gardner, J. F., Van Jaarsveld, R. H., De Lange, I. M., Van Der Smagt, J. J., Wilson, G. N., Dubbs, H., Goldberg, E. M., Zitano, L., Bupp, C., Martinez, J., Srour, M., Accogli, A., Alhakeem, A., Meltzer, M., Gropman, A., Brewer, C., Caswell, R. C., Montgomery, T., . . . Mefford, H. C. (2021). Variants in GNAI1 cause a syndrome associated with variable features including developmental delay, seizures, and hypotonia. Genetics in Medicine, 23(5), 881–887. 10.1038/s41436-020-01076-8

Muntean, B. S., Masuho, I., Dao, M., Sutton, L. P., Zucca, S., Iwamoto, H., Patil, D. N., Wang, D., Birnbaumer, L., Blakely, R. D., Grill, B., & Martemyanov, K. A. (2021). Gαo is a major determinant of cAMP signaling in the pathophysiology of movement disorders. Cell Reports, 34(5), 108718. 10.1016/j.celrep.2021.108718

Nechipurenko, I. V., Olivier-Mason, A., Kazatskaya, A., Kennedy, J., McLachlan, I. G., Heiman, M. G., Blacque, O. E., & Sengupta, P. (2016). A Conserved Role for Girdin in Basal Body Positioning and Ciliogenesis. Dev Cell, 38(5), 493–506. 10.1016/j.devcel.2016.07.013

Noel, J. P., Hamm, H. E., & Sigler, P. B. (1993). The 2.2 A crystal structure of transducin-alpha complexed with GTP gamma S. Nature, 366(6456), 654–663. 10.1038/366654a0

Oldham, W. M., Van Eps, N., Preininger, A. M., Hubbell, W. L., & Hamm, H. E. (2006). Mechanism of the receptor-catalyzed activation of heterotrimeric G proteins. Nat Struct Mol Biol, 13(9), 772–777. 10.1038/nsmb1129

Ott, C. M., Torres, R., Kuan, T. S., Kuan, A., Buchanan, J., Elabbady, L., Seshamani, S., Bodor, A. L., Collman, F., Bock, D. D., Lee, W. C., da Costa, N. M., & Lippincott-Schwartz, J. (2024). Ultrastructural differences impact cilia shape and external exposure across cell classes in the visual cortex. Curr Biol, 34(11), 2418–2433 e2414. 10.1016/j.cub.2024.04.043

Perkins, L. A., Hedgecock, E. M., Thomson, J. N., & Culotti, J. G. (1986). Mutant sensory cilia in the nematode Caenorhabditis elegans. Dev Biol, 117(2), 456–487. 10.1016/0012-1606(86)90314-3

Philbrook, A., O’Donnell, M. P., Grunenkovaite, L., & Sengupta, P. (2024). Cilia structure and intraflagellar transport differentially regulate sensory response dynamics within and between C. elegans chemosensory neurons. PLOS Biology, 22(11), e3002892. 10.1371/journal.pbio.3002892

Pierce, K. L., Premont, R. T., & Lefkowitz, R. J. (2002). Seven-transmembrane receptors. Nat Rev Mol Cell Biol, 3(9), 639–650. 10.1038/nrm908

Pineda, V. V., Athos, J. I., Wang, H., Celver, J., Ippolito, D., Boulay, G., Birnbaumer, L., & Storm, D. R. (2004). Removal of G(ialpha1) constraints on adenylyl cyclase in the hippocampus enhances LTP and impairs memory formation. Neuron, 41(1), 153–163. 10.1016/s0896-6273(03)00813-4

Posner, B. A., Mixon, M. B., Wall, M. A., Sprang, S. R., & Gilman, A. G. (1998). The A326S mutant of Gialpha1 as an approximation of the receptor-bound state. J Biol Chem, 273(34), 21752–21758. 10.1074/jbc.273.34.21752

Reiter, J. F., & Leroux, M. R. (2017). Genes and molecular pathways underpinning ciliopathies. Nature Reviews Molecular Cell Biology, 18(9), 533–547. 10.1038/nrm.2017.60

Roayaie, K., Crump, J. G., Sagasti, A., & Bargmann, C. I. (1998). The Gα Protein ODR-3 Mediates Olfactory and Nociceptive Function and Controls Cilium Morphogenesis in C. elegans Olfactory Neurons. Neuron, 20(1), 55–67. 10.1016/s0896-6273(00)80434-1

Seven, A. B., Hilger, D., Papasergi-Scott, M. M., Zhang, L., Qu, Q., Kobilka, B. K., Tall, G. G., & Skiniotis, G. (2020). Structures of Gα Proteins in Complex with Their Chaperone Reveal Quality Control Mechanisms. Cell Reports, 30(11), 3699–3709.e3696. 10.1016/j.celrep.2020.02.086

Sheu, S. H., Upadhyayula, S., Dupuy, V., Pang, S., Deng, F., Wan, J., Walpita, D., Pasolli, H. A., Houser, J., Sanchez-Martinez, S., Brauchi, S. E., Banala, S., Freeman, M., Xu, C. S., Kirchhausen, T., Hess, H. F., Lavis, L., Li, Y., Chaumont-Dubel, S., & Clapham, D. E. (2022). A serotonergic axon-cilium synapse drives nuclear signaling to alter chromatin accessibility. Cell, 185(18), 3390–3407 e3318. 10.1016/j.cell.2022.07.026

Shyu, Y. J., Hiatt, S. M., Duren, H. M., Ellis, R. E., Kerppola, T. K., & Hu, C.-D. (2008). Visualization of protein interactions in living Caenorhabditis elegans using bimolecular fluorescence complementation analysis. Nature Protocols, 3(4), 588–596. 10.1038/nprot.2008.16

Singh, J., Wen, X., & Scales, S. J. (2015). The Orphan G Protein-coupled Receptor Gpr175 (Tpra40) Enhances Hedgehog Signaling by Modulating cAMP Levels. J Biol Chem, 290(49), 29663–29675. 10.1074/jbc.M115.665810

Sprang, S. R. (1997). G protein mechanisms: insights from structural analysis. Annu Rev Biochem, 66, 639–678. 10.1146/annurev.biochem.66.1.639

Stoufflet, J., & Caille, I. (2022). The Primary Cilium and Neuronal Migration. Cells, 11(21). 10.3390/cells11213384

Suciu, S. K., & Caspary, T. (2021). Cilia, neural development and disease. Semin Cell Dev Biol, 110, 34–42. 10.1016/j.semcdb.2020.07.014

Sun, D., Flock, T., Deupi, X., Maeda, S., Matkovic, M., Mendieta, S., Mayer, D., Dawson, R., Schertler, G. F. X., Madan Babu, M., & Veprintsev, D. B. (2015). Probing Galphai1 protein activation at single-amino acid resolution. Nat Struct Mol Biol, 22(9), 686–694. 10.1038/nsmb.3070

Valente, E. M., Rosti, R. O., Gibbs, E., & Gleeson, J. G. (2014). Primary cilia in neurodevelopmental disorders. Nat Rev Neurol, 10(1), 27–36. 10.1038/nrneurol.2013.247

Varadi, M., Anyango, S., Deshpande, M., Nair, S., Natassia, C., Yordanova, G., Yuan, D., Stroe, O., Wood, G., Laydon, A., Zidek, A., Green, T., Tunyasuvunakool, K., Petersen, S., Jumper, J., Clancy, E., Green, R., Vora, A., Lutfi, M., . . . Velankar, S. (2022). AlphaFold Protein Structure Database: massively expanding the structural coverage of protein-sequence space with high-accuracy models. Nucleic Acids Res, 50(D1), D439–D444. 10.1093/nar/gkab1061

Varadi, M., Bertoni, D., Magana, P., Paramval, U., Pidruchna, I., Radhakrishnan, M., Tsenkov, M., Nair, S., Mirdita, M., Yeo, J., Kovalevskiy, O., Tunyasuvunakool, K., Laydon, A., Zidek, A., Tomlinson, H., Hariharan, D., Abrahamson, J., Green, T., Jumper, J., . . . Velankar, S. (2024). AlphaFold Protein Structure Database in 2024: providing structure coverage for over 214 million protein sequences. Nucleic Acids Res, 52(D1), D368–D375. 10.1093/nar/gkad1011

Wang, D., Dao, M., Muntean, B. S., Giles, A. C., Martemyanov, K. A., & Grill, B. (2022). Genetic modeling of GNAO1 disorder delineates mechanisms of Gαo dysfunction. Human Molecular Genetics, 31(4), 510–522. 10.1093/hmg/ddab235

Wettschureck, N., & Offermanns, S. (2005). Mammalian G proteins and their cell type specific functions. Physiol Rev, 85(4), 1159–1204. 10.1152/physrev.00003.2005

Wong, W. R., Brugman, K. I., Maher, S., Oh, J. Y., Howe, K., Kato, M., & Sternberg, P. W. (2019). Autism-associated missense genetic variants impact locomotion and neurodevelopment in Caenorhabditis elegans. Hum Mol Genet, 28(13), 2271–2281. 10.1093/hmg/ddz051

Wright, K. J., Baye, L. M., Olivier-Mason, A., Mukhopadhyay, S., Sang, L., Kwong, M., Wang, W., Pretorius, P. R., Sheffield, V. C., Sengupta, P., Slusarski, D. C., & Jackson, P. K. (2011). An ARL3-UNC119-RP2 GTPase cycle targets myristoylated NPHP3 to the primary cilium. Genes Dev, 25(22), 2347–2360. 10.1101/gad.173443.111

Wu, J. Y., Cho, S. J., Descant, K., Li, P. H., Shapson-Coe, A., Januszewski, M., Berger, D. R., Meyer, C., Casingal, C., Huda, A., Liu, J., Ghashghaei, T., Brenman, M., Jiang, M., Scarborough, J., Pope, A., Jain, V., Stein, J. L., Guo, J., . . . Anton, E. S. (2024). Mapping of neuronal and glial primary cilia contactome and connectome in the human cerebral cortex. Neuron, 112(1), 41–55 e43. 10.1016/j.neuron.2023.09.032

Xing, S., Wallmeroth, N., Berendzen, K. W., & Grefen, C. (2016). Techniques for the analysis of protein-protein interactions in vivo. Plant Physiology, pp.00470.02016. 10.1104/pp.16.00470

Yoshida, K., Hirotsu, T., Tagawa, T., Oda, S., Wakabayashi, T., Iino, Y., & Ishihara, T. (2012). Odour concentration-dependent olfactory preference change in C. elegans. Nat Commun, 3, 739. 10.1038/ncomms1750

Zeng, B., Mou, T.-C., Doukov, T. I., Steiner, A., Yu, W., Papasergi-Scott, M., Tall, G. G., Hagn, F., & Sprang, S. R. (2019). Structure, Function, and Dynamics of the Gα Binding Domain of Ric-8A. Structure, 27(7), 1137-1147.e1135. 10.1016/j.str.2019.04.013

Zhang, H., Constantine, R., Vorobiev, S., Chen, Y., Seetharaman, J., Huang, Y. J., Xiao, R., Montelione, G. T., Gerstner, C. D., Davis, M. W., Inana, G., Whitby, F. G., Jorgensen, E. M., Hill, C. P., Tong, L., & Baehr, W. (2011). UNC119 is required for G protein trafficking in sensory neurons. Nat Neurosci, 14(7), 874–880. 10.1038/nn.2835

