## Supplementary material for "Functional classification of *GNAI1* disorder variants in *C. elegans* uncovers conserved and cell-specific mechanisms of dysfunction"

Supplementary information

### MATERIALS AND METHODS

#### Molecular Biology

*BiFC constructs*: The cDNA sequence encoding the C-terminal fragment of Venus was amplified from pCe-BiFC-VC155 (Addgene) and subcloned into *C. elegans* expression vector containing *bbs-8*p*::ric-8* sequence using standard restriction enzyme cloning. The cDNA sequence encoding the N-terminal fragment of Venus was amplified from pCe-BiFC-VN173 (Addgene) and subcloned into pMC10 *C. elegans* expression vector containing *ceh-36∆*p*::odr-3*$^{A328P}$ sequence using NEBuilder HiFi assembly (NEB). The resulting plasmids were verified by whole-plasmid sequencing (Plasmidsaurus). The *unc-119* cDNA was amplified from a mixed-stage N2 cDNA library and assembled into *C. elegans* expression vector to generate *bbs-8*p*::VC155::unc-119* plasmid using NEBuilder HiFi DNA assembly (NEB).

#### Fluorescence lifetime imaging microscopy (FLIM)

FLIM of AWC sensory endings was carried out on Alba v5 instrument (ISS) using a two-photon titanium-sapphire laser and a Nikon Eclipse Ti-U inverted microscope. FastFLIM mode in VistaVision software (ISS) was used for data acquisition. Atto 425 fluorescent dye ($\tau$ = 3.61 ns at 80MHz) was used to calibrate the samples' lifetimes. All images were acquired at identical settings and on at least two independent days.

#### Bimolecular fluorescence complementation (BiFC)

One-day-old transgenic worms expressing complementary pairs of BiFC constructs were imaged on an inverted Nikon Ti-E microscope with Yokogawa CSU-X1 spinning disk confocal head using a 60X NA 1.40 oil immersion objective. Complete z-stacks of the anterior end of transgenic animals were acquired at 0.25-µm intervals with an ORCA-Fusion BT Digital CMOS camera (Hamamatsu) in MetaMorph 7 software (Molecular devices).

#### Measurement of ASH cilia length

To measure ASH cilium length, a line was drawn from the base of ASH cilium to the tip using the segmented line tool in Fiji/Image J (National Institute of Health, Bethesda, MD).

### FIGURE LEGENDS

**Supplementary Fig. 1.** *GNAI1* knockdown in RPE-1 cells impairs ciliogenesis. a) Quantification of ciliation in RPE-1 cells transfected with siControl or si*GNAI1* #2 and stained with anti-ARL13B antibody to visualize cilia. Each data point represents one replicate. Data for siControl are repeated from Fig. 1c. Total number of cells: siControl (n=858), si*GNAI1* #2 (n=400). Means ± SD are indicated by shaded and vertical bars, respectively. *** indicates different from control at $p<0.001$ (Fisher's exact test). b) Relative levels of *GNAI1* mRNA in RPE-1 cells treated with the indicated siRNAs. Data for siControl are repeated from Fig. 1e. Summary data represent > 3 replicates. Error bars are SEM. * indicates different from control at $p<0.05$ (Mann-Whitney test). c) Representative images of fixed hTERT RPE-1 cells transfected with the indicated siRNAs and stained with anti-acetylated tubulin antibody and DAPI. Scale: 10 μm. d) Quantification of ciliation in RPE-1 cells transfected with the indicated siRNAs and stained with anti-acetylated tubulin antibody. Each data point represents one replicate. Total number of cells: siControl (n=719), si*GNAI1* #1 (n=273). Means ± SD are indicated by shaded and vertical bars, respectively. *** indicates different from control at $p<0.001$ (Fisher's exact test).

**Supplementary Fig. 2.** *GNAI1* disorder variants alter identical residues in conserved motifs of ODR-3. Sequence alignment of ODR-3 and human Gαi1 showing patient and orthologous ODR-3 mutations (orange boxes) that were selected for analysis.

**Supplementary Fig. 3.** The *A328P* variant disrupts cilia localization of the endogenous ODR-3. Representative images of ODR-3$^{WT}$::split-wrmScarlet and ODR-3$^{A328P}$::split-wrmScarlet. Purple arrowheads mark cilia base. White arrow points to autofluorescence in the pharynx. d: dendrite. Anterior is at left. Scale: 10 $\mu$m. Total number of examined animals: ODR-3$^{WT}$ (n=25) and ODR-3$^{A328P}$ (n=30).

**Supplementary Fig. 4.** crRNA target sequences and CRISPR-engineered mutations in the *odr-3* gene. a – f) CRISPR design for a) *T48I*, b) *M88V*, c) *D175V*, d) *I321T*, e) *A328P*, and f) *V334E* variants. Altered codons are highlighted in pink with the corresponding amino acids shown above and/or below codon sequences. Missense mutations are shown in pink (pink asterisks). Silent mutations introduced in the donor sequence to prevent Cas9-mediated re-cutting are shown in dark blue (dark blue asterisks). crRNA target sequences and PAM sites are in light blue and orange, respectively. All donor templates also contained 36-40-base pair homology arms (partial homology-arm sequences are shown for *T48I*, *D175V*, and *I321T* variants due to space limitation).

**Supplementary Fig. 5.** DNA sequencing verification of the CRISPR-engineered orthologous patient variants. a– f) Raw traces of Sanger DNA sequencing results for a) *T48I*, b) *M88V*, c) *D175V*, d) *I321T*, e) *A328P*, and f) *V334E* mutations in the *odr-3* gene. Blue asterisks mark silent mutations that were introduced to prevent Cas9-mediated re-cutting.

**Supplementary Fig. 6.** Wild-type and A328P ODR-3 variants physically interact with RIC-8. a) Representative images and b) quantification of BiFC in transgenic animals overexpressing indicated constructs in AWC neurons. Purple arrowheads mark cilia base. White dashed lines mark boundaries of the worm nose. Anterior is at left. Scale: 10 $\mu$m. VN and VC: N- and C-terminal fragments of Venus, respectively. The number of analyzed animals per genotype is indicated above corresponding bars on the graph. ** and *** indicate different from the negative control at $p<0.01$ and $p<0.001$, respectively (Fisher's exact test). c) Quantification of lifetimes of

RIC-8::GFP in the presence of the indicated TagRFP-tagged ODR-3 variants. All constructs were expressed under the AWC-specific *ceh-36*Δp regulatory sequences. Each point corresponds to a single AWC neuron. Means ± SD are indicated by shaded and vertical bars, respectively. The number of analyzed animals per genotype is indicated above corresponding bars on the graph. ** and *** indicate different from RIC-8::GFP alone at $p<0.01$ and $p<0.001$, respectively (Kruskal-Wallis with Dunn's multiple comparisons test). d) Representative phasor plots for distal AWC endings in animals of the indicated genotypes. Each point represents an intensity pixel value from the imaged AWC sensory ending.

**Supplementary Fig. 7.** Wild-type and A328P ODR-3 variants associate with UNC-119. a) Representative images and b) quantification of BiFC in transgenic animals overexpressing indicated constructs in AWC neurons. Purple arrowheads mark cilia base. White dashed lines mark boundaries of the worm nose. Anterior is at left. Scale: 10 μm. VN and VC: N- and C-terminal fragments of Venus, respectively. The number of analyzed animals per genotype is indicated above corresponding bars on the graph. * and *** indicate different from the negative control at $p<0.05$ and $p<0.001$, respectively (Fisher's exact test).

**Supplementary Fig. 8.** ASH cilium length is normal in *odr-3(lf)* mutants. a) Representative images and b) quantification of ASH cilium length in animals of the indicated genotypes. Anterior is at left. Scale: 5 μm. Total number of cilia: WT (n=36), *odr-3(lf)* (n=29). Means ± SD are indicated by shaded and vertical bars, respectively. ns: not significant (unpaired t test with Welch's correction).

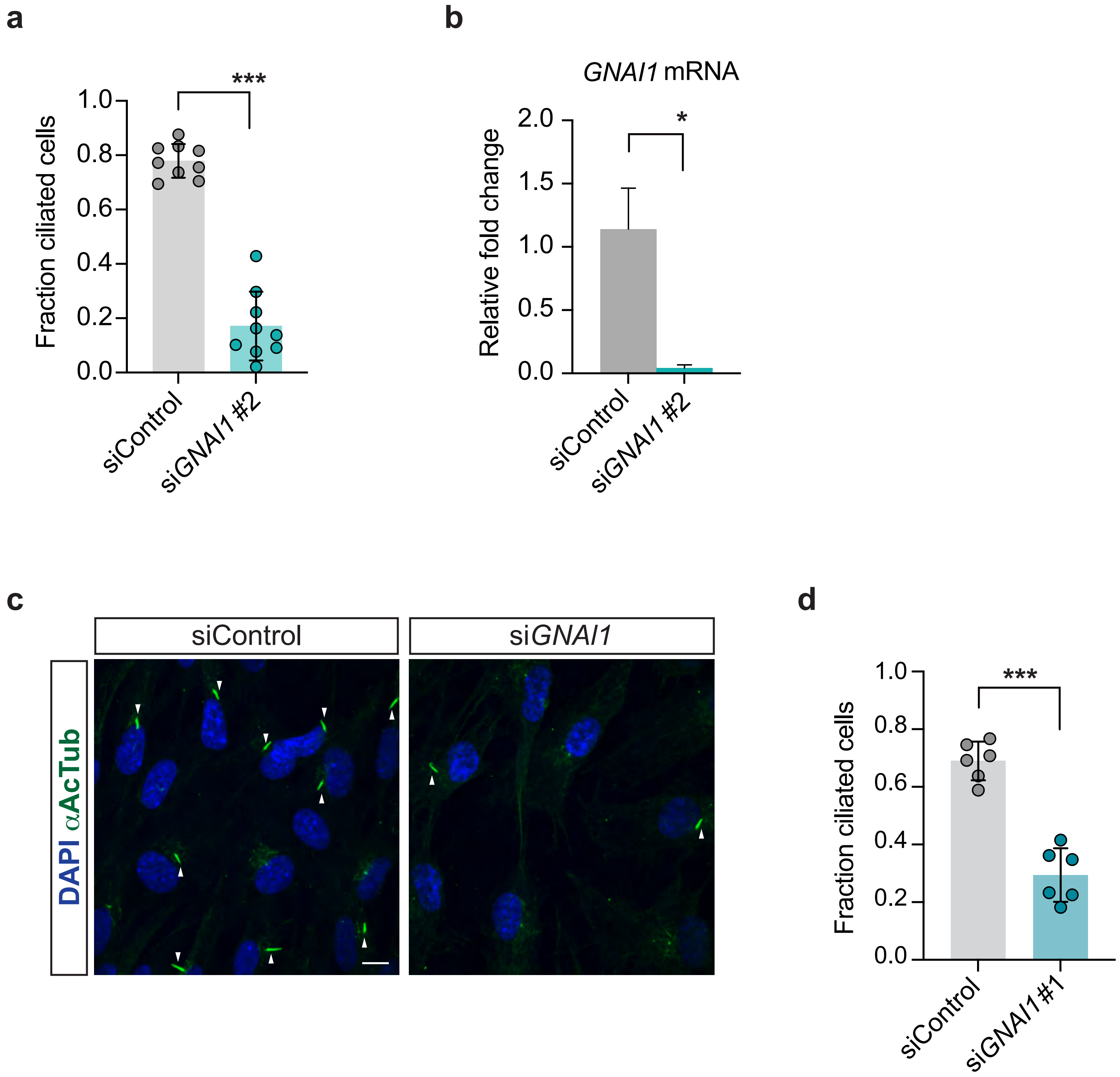


Salama *et al.* **Supplementary Figure 1**

```
                                                       T48I
ODR-3   MGSCQSNENSEGNARNKEIEKQLNADKRAGSSIVKLLLLGAGECGKSTVLKQMQILHSNG 60
Gαi1    MGCTLSAEDKAAVERSKMIDRNLREDGEKAAREVKLLLLGAGESGKSTIVKQMKIIHEAG 60
        **.  * *:. .  *.* *:::*. * . .:  **********.****::***:*:*. *

                                     M88V
ODR-3   FTEEEVNEKRAIVYNNTVSAMCTILRAMDGVLHLPLENGQKEAEKAIVMKVQENGEEGEA 120
Gαi1    YSEEECKQYKAVVYSNTIQSIIAIIRAMGRL-KIDFGDSARADDAR-QLFVLAGAAEEGF 118
        ::*** :: :*:**.**:.:: :*:***. : :: : :. :  :    : *  .. *

                                                           D175V
ODR-3   LTEEVSKAIQSLWADPGVKKAFEMRSEYQLPDSAKYFLDNCQRISEPGYRPNDQDILYSR 180
Gαi1    MTAELAGVIKRLWKDSGVQACFNRSREYQLNDSAAYYLNDLDRIAQPNYIPTQQDVLRTR 178
        :* *:: .*: ** * **: .*:   **** *** *:*:: :**::*.* *.:**:* :*
                                                           D173V

ODR-3   VATTGVVEVKFKIKELDFRVFDVGGQRSERRKWIHCFDNVESIIFITAISEYDQVLFEDE 240
Gαi1    VKTTGIVETHFTFKDLHFKMFDVGGQRSERKKWIHCFEGVTAIIFCVALSDYDLVLAEDE 238
        * ***:**.:*.:*:*.*::**********:******:.* :*** .*:*:** ** ***

                                        K272R
ODR-3   TTNRMIESMQLFNSICNSTWFLSTAMILFMNKKDLFMEKIQRVNITTAFPDYEGGQNYEE 300
Gαi1    EMNRMHESMKLFDSICNNKWFTDTSIILFLNKKDLFEEKIKKSPLTICYPEYAGSNTYEE 298
          *** ***:**:****..** .*::***:****** ***::  :* .:*:* *.:.***
                                        K270R
                   I321T   A328P  V334E
ODR-3   AVSFIKQKFAELNLNPDKKTIYMHETCATDTNQVQLVISSVIDTIIQKNLQKAGMM     356
Gαi1    AAAYIQCQFEDLNKRKDTKEIYTHFTCATDTKNVQFVFDAVTDVIIKNNLKDCGLF     354
        *.::*: :* :** .  *.* ** * ******::**:*:.:* *.**:**:..*::
                   I319T   A326P  V332E
```

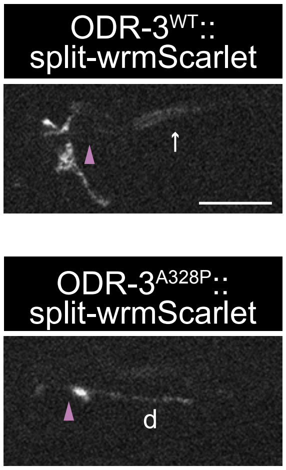
ODR-3WT::
split-wrmScarlet
ODR-3A328P::
split-wrmScarlet
d

**A**

**WT** (T)

CTGGAAGTAGTATTGTTAAGCTTTTGCTACTTGGAGCAGGAGAATGTGGAAAATCTACAGTTCTGAAACAGATGCAgtat

***T48I*** (I)

CTGGAAGTAGTATTGTTAAGCTTTTGCTACTTGGAGC**T**GG**G**GAATGTGGAAAATCTA**T**AGTTCTGAAACAGATGCAgtat

**B**

**WT** (M)

CAATAATACTGTTTCTGCAATGTGCACAATTTTAAGAGCAATGGACGGAGTTCTGCATTTACCGTTGGAAAATGGACAAAAAG

***M88V*** (V)

CAATAATACTGTTTCTGCAATGTGCAC**T**ATTTTAAGAGC**TG**TGGACGGAGTTCTGCATTTACCGTTGGAAAATGGACAAAAAG

**C**

**WT** (D)

TAATTGTCAGCGTATATCTGAACCTGGGTATCGGCCAAACGATCAAGATATTCTTTATTCTCGAGTGGCAACAACTGGTGTCGT

***D175V*** (V)

TAATTGTCAGCGTATATCTGAACCTGGGTATCGGCCAAA**T**GATCAAG**T**TAT**C**CTTTATTCTCGAGTGGCAACAACTGGTGTCGT

**D**

**WT** (I)

ATCCTGATAAAAAGACAATTTATATGCATGAAACTTGCGCTACAGACACTAATCAGgtaggttattggtttattggaattttaattctgaactgatcttca

***I321T*** (T)

ATCCTGATAAAAAGACAA**C**TTATATGCATGAAACTTGCGCTACAGACACTAATCAGgtaggttatt**cc**tttattggaattttaattctgaactgatcttca

**E**

**WT** (A)

GAATCCTGATAAAAAGACAATTTATATGCATGAAACTTGCGCTACAGACACTAATCAGgtaggttattggtttattggaattttaattctgaactgatcttcaa

***A328P*** (P)

GAATCCTGATAAAAAGACAATTTATATGCATGAAACTTGC**C**CTACAGACACTAATCAGgtaggttatt**cc**tttattggaattttaattctgaactgatcttcaa

**F**

**WT** (V)

tctgaactgatcttcaaaaaaaaaacttaattcccaattttcagGTACAACTTGTCATTTCGAGTGTTATTGATACAATTATTCAG

***V334E*** (E)

tctgaactgatcttcaaaaaaaaaacttaattc**g**caattttcagG**A**ACAACTTGT**T**ATTTCGAGTGTTATTGATACAATTATTCAG

Salama *et al.* **Supplementary Figure 4**

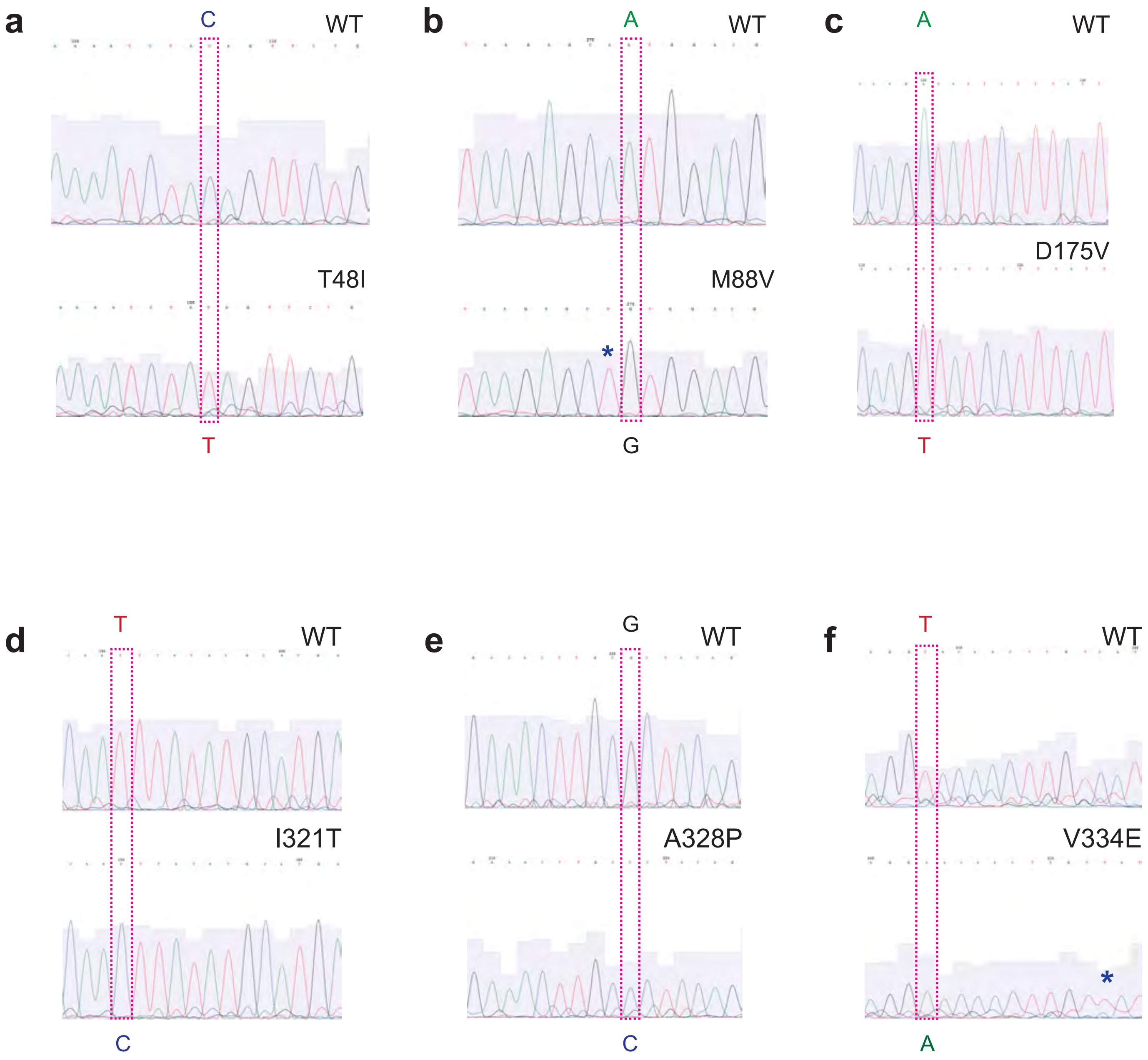
a
C
WT
T48I
T
b
A
WT
M88V
*
G
c
A
WT
D175V
T
d
T
WT
I321T
C
e
G
WT
A328P
C
f
T
WT
V334E
*
A

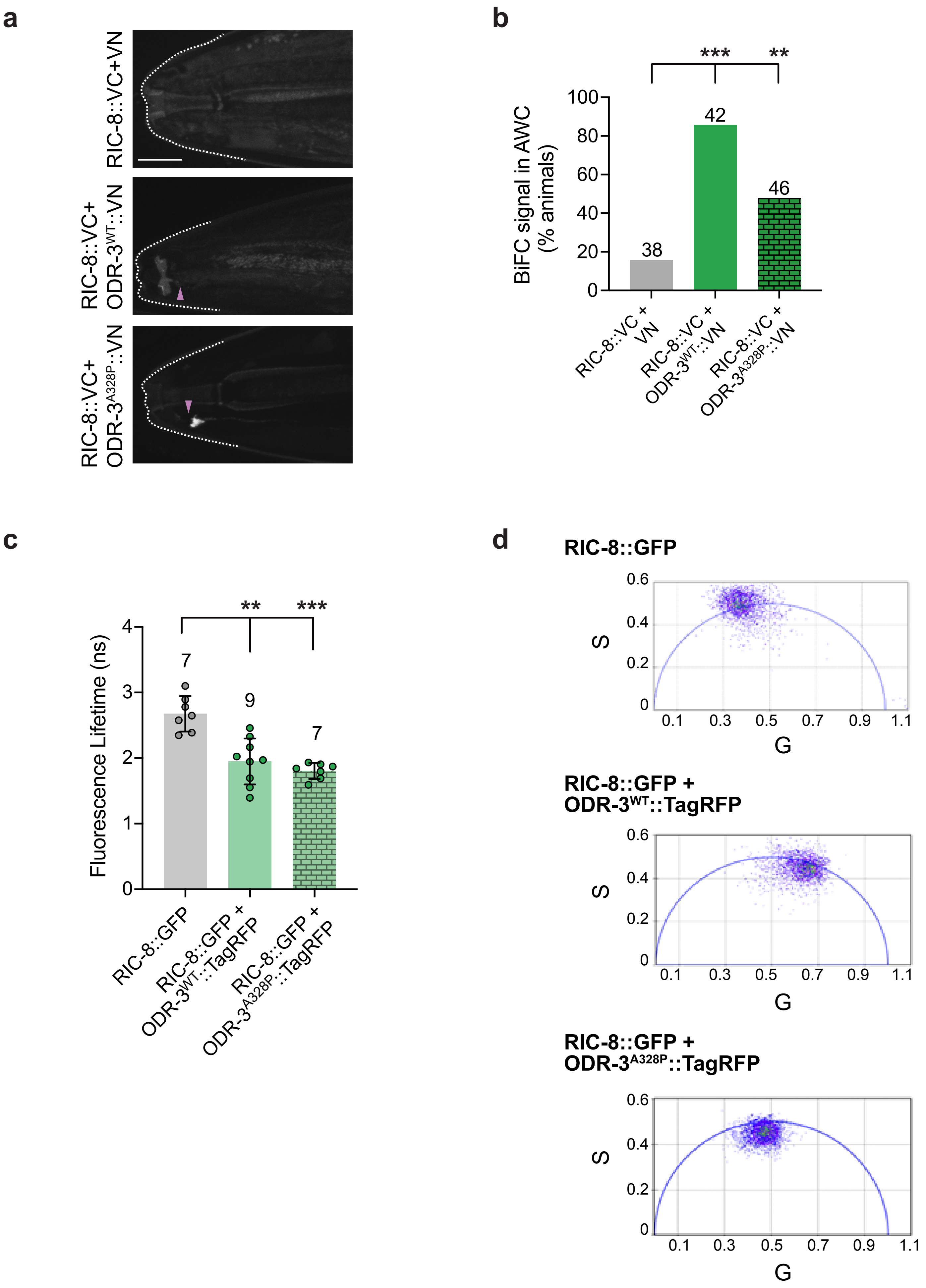
a
RIC-8::VC+VN
RIC-8::VC+ ODR-3WT::VN
RIC-8::VC+ ODR-3A328P::VN
b
BiFC signal in AWC (% animals)
***
**
100
80
60
40
20
0
38
42
46
RIC-8::VC + VN
RIC-8::VC + ODR-3WT::VN
RIC-8::VC + ODR-3A328P::VN
c
Fluorescence Lifetime (ns)
**
***
4
3
2
1
0
7
9
7
RIC-8::GFP
RIC-8::GFP + ODR-3WT::TagRFP
RIC-8::GFP + ODR-3A328P::TagRFP
d
RIC-8::GFP
RIC-8::GFP + ODR-3WT::TagRFP
RIC-8::GFP + ODR-3A328P::TagRFP
S
G
0.6
0.4
0.2
0
0.1
0.3
0.5
0.7
0.9
1.1

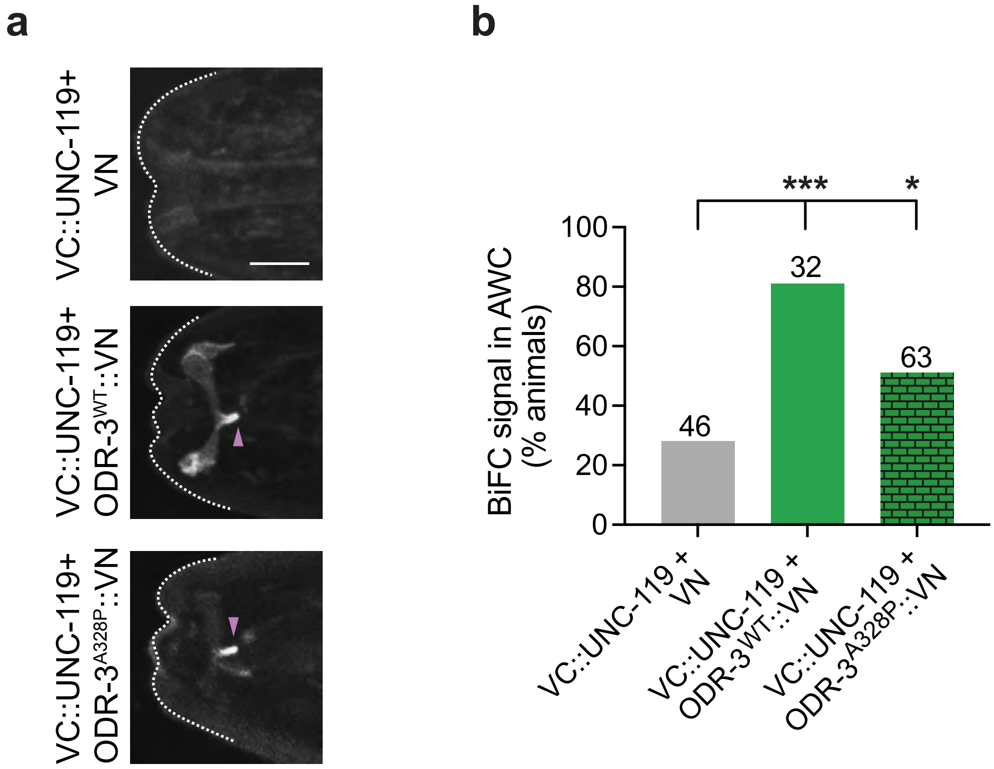
a
VC::UNC-119+ VN
VC::UNC-119+ ODR-3WT::VN
VC::UNC-119+ ODR-3A328P::VN
b
***
*
BiFC signal in AWC (% animals)
100
80
60
40
20
0
46
32
63
VC::UNC-119 + VN
VC::UNC-119 + ODR-3WT::VN
VC::UNC-119 + ODR-3A328P::VN

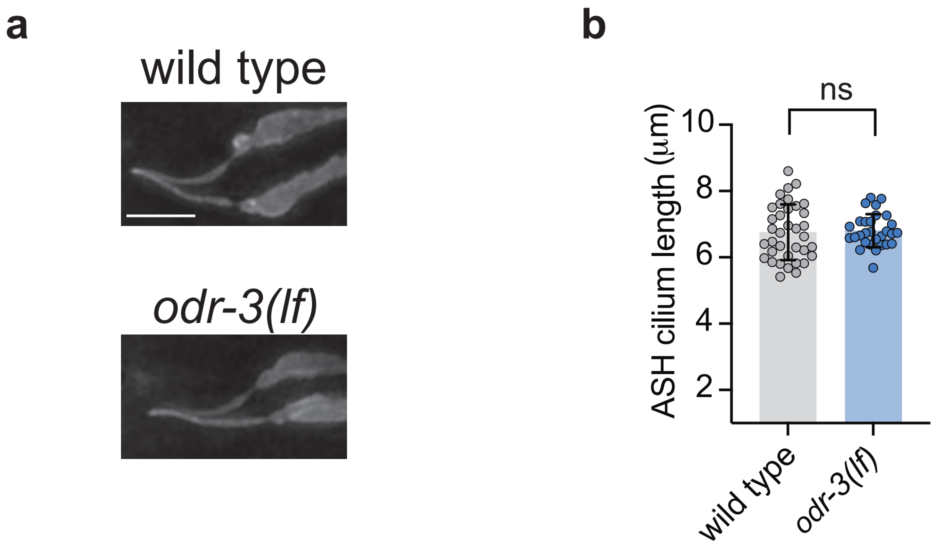

a
wild type
odr-3(lf)
b
ns
10
8
6
4
2
ASH cilium length (μm)
wild type
odr-3(lf)

**Supplementary Table 1**: List of *C. elegans* strains used in this work

| Strain | Genotype | Source |
|---|---|---|
| NWM868 | *odr-3(nch013[odr-3::wrmScarlet$_{11}$])* V*; nchEx021[ceh-36△*p*::wrmScarlet$_{1-10}$, unc-122△*p*::gfp]* | This work |
| NWM447 | *oyIs50[ceh-36*p*::gfp]* IV; *nchEx025[ceh-36△*p*::odr-3::tagrfp, unc-122△*p*::dsRed]* | This work |
| NWM456 | *oyIs50[ceh-36*p*::gfp]* IV; *nchEx026[ceh-36△*p*::odr-3$^{I321T}$::tagrfp, unc-122△*p*::dsRed]* | This work |
| NWM457 | *oyIs50[ceh-36*p*::gfp]* IV; *nchEx027[ceh-36△*p*::odr-3$^{I321T}$::tagrfp, unc-122△*p*::dsRed]* | This work |
| NWM514 | *oyIs50[ceh-36*p*::gfp]* IV; *nchEx033[ceh-36△*p*::odr-3$^{V334E}$::tagrfp, unc-122△*p*::dsRed]* | This work |
| NWM515 | *oyIs50[ceh-36*p*::gfp]* IV; *nchEx034[ceh-36△*p*::odr-3$^{V334E}$::tagrfp, unc-122△*p*::dsRed]* | This work |
| NWM576 | *oyIs50[ceh-36*p*::gfp]* IV; *nchEx019[ceh-36△*p*::odr-3$^{T48I}$::tagrfp, unc-122△*p::*dsRed]* | This work |
| NWM577 | *oyIs50[ceh-36*p*::gfp]* IV; *nchEx020[ceh-36△*p*::odr-3$^{T48I}$::tagrfp, unc-122△*p::*dsRed]* | This work |
| NWM472 | *oyIs50[ceh-36*p*::gfp]* IV; *nchEx022[ceh-36△*p*::odr-3$^{M88V}$::tagrfp, unc-122△*p*::dsRed]* | This work |
| NWM474 | *oyIs50[ceh-36*p*::gfp]* IV; *nchEx023[ceh-36△*p*::odr-3$^{M88V}$::tagrfp, unc-122△*p*::dsRed]* | This work |
| NWM445 | *oyIs50[ceh-36*p*::gfp]* IV; *nchEx035 [ceh-36△*p*::odr-3$^{D175V}$::tagrfp, unc-122△*p*::dsRed]* | This work |
| NWM446 | *oyIs50[ceh-36*p*::gfp]* IV; *nchEx032 [ceh-36△*p*::odr-3$^{D175V}$::tagrfp, unc-122△*p*::dsRed]* | This work |
| NWM505 | *oyIs50[ceh-36*p*::gfp]* IV; *nchEx030[ceh-36△*p*::odr-3$^{K272R}$::tagrfp, unc-122△*p*::dsRed]* | This work |
| NWM506 | *oyIs50[ceh-36*p*::gfp]* IV; *nchEx031[ceh-36△*p*::odr-3$^{K272R}$::tagrfp, unc-122△*p*::dsRed]* | This work |
| NWM494 | *oyIs50[ceh-36*p*::gfp]* IV; *nchEx028[ceh-36△*p*::odr-3$^{A328P}$::tagrfp, unc-122△*p*::dsRed]* | This work |
| NWM500 | *oyIs50[ceh-36*p*::gfp]* IV; *nchEx029[ceh-36△*p*::odr-3$^{A328P}$::tagrfp, unc-122△*p*::dsRed]* | This work |
| NWM869 | *odr-3(nch027[odr-3[A328P]::wrmScarlet$_{11}$])* V*; nchEx021[ceh-36△*p*::wrmScarlet$_{1-10}$, unc-122△*p*::gfp]* | This work |
| NWM862 | *mks-5(tm3100)* II*; oyIs50[ceh-36*p*::gfp]* IV*; nchEx028[ceh-36△*p*::odr-3$^{A328P}$::tagrfp, unc-122△*p*::dsRed]* | This work |
| NWM861 | *mks-5(tm3100)* II*; oyIs50[ceh-36*p*::gfp]* IV*; nchEx025[ceh-36△*p*::odr-3::tagrfp, unc-122△*p*::dsRed]* | This work |
| PY3453 | *oyIs50[ceh-36*p*::gfp]* IV | (Kim, Kim et al. 2010) |
| NWM185 | *oyIs50[ceh-36*p*::gfp]* IV; *odr-3(n1605)* V | (Campagna, McMahon et al. 2023) |
| NWM448 | *oyIs50[ceh-36*p*::gfp]* IV*; odr-3(nch004[M88V])* V | This work |
| NWM502 | *oyIs50[ceh-36*p*::gfp]* IV*; odr-3(nch005[I321T])* V | This work |
| NWM563 | *oyIs50[ceh-36*p*::gfp]* IV*; odr-3(nch012[T48I])* V | This work |

| | | |
|---|---|---|
| NWM545 | *oyIs50[ceh-36*p*::gfp]* IV*; odr-3(nch011[D175V])* V | This work |
| NWM544 | *oyIs50[ceh-36*p*::gfp]* IV; *odr-3(nch009[A328P])* V | This work |
| NWM540 | *oyIs50[ceh-36*p*::gfp]* IV; *odr-3(nch0010[V334E])* V | This work |
| NWM710 | *oyIs50[ceh-36*p*::gfp]* IV; *odr-3* $^{A328P}$*(nch009)* V*; nchEx025[ceh-36△*p*::odr-3::tagrfp, unc-122△*p*::dsRed]* | This work |
| NWM652 /653 | *oyIs-50[ceh-36*p*::gfp]* IV; *odr-3(n1605)* V*; nchEx031[ceh-36△p::odr-3*$^{K272R}$*::tagrfp, unc-122△*p*::dsRed]* | This work |
| NWM864 | *nchEx042(bbs-8*p*::ric-8::vc155, ceh-36△*p*::vn173, unc-122△*p*::dsRed]* | This work |
| NWM865 | *nchEx043(bbs-8*p*::ric-8::vc155, ceh-36△*p*::vn173, unc-122△*p*::dsRed]* | This work |
| NWM765 | *nchEx038(bbs-8*p*::ric-8::vc155, ceh-36△*p*::odr-3::vn173, unc-122△*p*::dsRed]* | This work |
| NWM767 | *nchEx039(bbs-8*p*::ric-8::vc155, ceh-36△*p*::odr-3::vn173, unc-122△*p*::dsRed]* | This work |
| NWM831 | *nchEx040(bbs-8*p*::ric-8::vc155, ceh-36△*p*::odr-3*$^{A328P}$*::vn173, unc-122△*p*::dsRed]* | This work |
| NWM832 | *nchEx041(bbs-8*p*::ric-8::vc155, ceh-36△*p*::odr-3*$^{A328P}$*::vn173, unc-122△*p*::dsRed]* | This work |
| NWM858 | *nchEx045(ceh-36△*p*::ric-8::gfp, unc-122△*p*::dsRed)* | This work |
| NWM859 | *nchEx046(ceh-36△*p*::ric-8::gfp, ceh-36△*p*::odr-3::tagrfp, unc-122△*p*::dsRed)* | This work |
| NWM860 | *nchEx047(ceh-36△*p*::ric-8::gfp, ceh-36△*p*::odr-3*$^{A328P}$*::tagrfp, unc-122△*p*::dsRed)* | This work |
| NWM737 | *nchEx048(bbs-8*p*::vc155::unc-119, ceh-36△*p*::vn173, unc-122△*p*::dsRed]* | This work |
| NWM738 | *nchEx049(bbs-8*p*::vc155::unc-119, ceh-36△*p*::vn173, unc-122△*p*::dsRed]* | This work |
| NWM699 | *nchEx050(bbs-8*p*::vc155::unc-119, ceh-36△*p*::odr-3::vn173, unc-122△*p*::dsRed]* | This work |
| NWM727 | *nchEx051(bbs-8*p*::vc155::unc-119, ceh-36△*p*::odr-3*$^{A328P}$*::vn173, unc-122△*p*::dsRed]* | This work |
| NWM728 | *nchEx052(bbs-8*p*::vc155::unc-119, ceh-36△*p*::odr-3*$^{A328P}$*::vn173, unc-122△*p*::dsRed]* | This work |
| NWM437 | *nchEx036[sra-6*p*::myr-gfp, sra-6*p*::mksr-2::tagrfp, unc-122△*p*::dsRed]* | This work |
| NWM863 | *odr-3(n1605)* V*, nchEx036[sra-6△*p*::myr-gfp, sra-6△*p*::mksr-2::tagrfp, unc-122△*p*::dsRed]* | This work |
| NWM709 | *nchEx037[sra-6*p*::myr-gfp, sra-6*p*::odr-3::tagrfp, unc-122△*p*::dsRed]* | This work |
| NWM649 | *nchEx044[sra-6*p*::myr-gfp, sra-6*p*::odr-3*$^{D175V}$*::tagrfp, unc-122△*p*::dsred]* | This work |
| NWM711 | *odr-3(nch011[D175V])* V; *nchEx037[sra-6*p*::myr-gfp, sra-6*p*::odr-3::tagrfp, unc-122△*p*::dsRed]* | This work |

### REFERENCES


Campagna, C. M., H. McMahon and I. Nechipurenko (2023). "The G protein alpha chaperone and guanine-nucleotide exchange factor RIC-8 regulates cilia morphogenesis in Caenorhabditis elegans sensory neurons." PLoS Genetics **19**(11): e1011015.

Kim, K., R. Kim and P. Sengupta (2010). "The HMX/NKX homeodomain protein MLS-2 specifies the identity of the AWC sensory neuron type via regulation of the ceh-36 Otx gene in C. elegans." Development **137**(6): 963-974.

**Supplementary Table 2:** List of plasmids used in this work

| Plasmid | Description | Source |
|---|---|---|
| NWM084 | *ceh-36Δ*p*::wrmScarlet$_{1-10}$* | This work |
| Co-injection marker | *unc-122Δ*p*::gfp* | (Miyabayashi et al., 1999) |
| NWM032 | *ceh-36Δ*p*::odr-3::tagrfp* | (Campagna et al., 2023) |
| Co-injection marker | *unc-122Δ*p*::dsRed* | (Miyabayashi et al., 1999) |
| NWM051 | *ceh-36Δ*p*::odr-3$^{I321T}$::tagrfp* | This work |
| NWM055 | *ceh-36Δ*p*::odr-3$^{V334E}$::tagrfp* | This work |
| NWM059 | *ceh-36Δ*p*::odr-3$^{T48I}$::tagrfp* | This work |
| NWM050 | *ceh-36Δ*p*::odr-3$^{M88V}$::tagrfp* | This work |
| NWM052 | *ceh-36Δ*p*::odr-3$^{D175V}$::tagrfp* | This work |
| NWM053 | *ceh-36Δ*p*::odr-3$^{K272R}$::tagrfp* | This work |
| NWM054 | *ceh-36Δ*p*::odr-3$^{A328P}$::tagrfp* | This work |
| NWM036 | *bbs-8*p*::ric-8::vc155* | This work |
| NWM082 | *ceh-36Δ*p*::vn173* | (Campagna et al., 2023) |
| NWM034 | *ceh-36Δ*p*::odr-3$^{WT}$::vn173* | (Campagna et al., 2023) |
| NWM095 | *ceh-36Δ*p*::odr-3$^{A328P}$::vn173* | This work |
| NWM043 | *ceh-36Δ*p*::ric-8::gfp* | This work |
| NWM116 | *bbs-8*p*::vc155::unc-119* | This work |
| NWM100 | *sra-6p::myrgfp* | This work |
| NWM044 | *sra-6p::mksr-2::tagrfp* | This work |
| NWM097 | *sra-6*p*::odr-3$^{WT}$::tagrfp* | This work |
| NWM099 | *sra-6*p*::odr-3$^{D175V}$::tagrfp* | This work |
| NWM049 | *GNAI1$^{WT}$::eGFP pcDNA3.1+* | This work |
| NWM065 | *GNAI1$^{D173V}$::eGFP pcDNA3.1+* | This work |
| NWM066 | *GNAI1$^{K270R}$::eGFP pcDNA3.1+* | This work |
| NWM068 | *GNAI1$^{A326P}$::eGFP pcDNA3.1+* | This work |

### REFERENCES


Campagna, C. M., McMahon, H., & Nechipurenko, I. (2023). The G protein alpha chaperone and guanine-nucleotide exchange factor RIC-8 regulates cilia morphogenesis in Caenorhabditis elegans sensory neurons. *PLoS Genetics*, *19*(11), e1011015. https://doi.org/10.1371/journal.pgen.1011015

Miyabayashi, T., Palfreyman, M. T., Sluder, A. E., Slack, F., & Sengupta, P. (1999). Expression and function of members of a divergent nuclear receptor family in Caenorhabditis elegans. *Dev Biol*, *215*(2), 314-331. https://doi.org/10.1006/dbio.1999.9470
